# Welcome to the big leaves: best practices for improving genome annotation in non-model plant genomes

**DOI:** 10.1101/2022.10.03.510643

**Authors:** Vidya S Vuruputoor, Daniel Monyak, Karl C. Fetter, Cynthia Webster, Akriti Bhattarai, Bikash Shrestha, Sumaira Zaman, Jeremy Bennett, Susan L. McEvoy, Madison Caballero, Jill L. Wegrzyn

## Abstract

**Premise of the study:** Robust standards to evaluate quality and completeness are lacking for eukaryotic structural genome annotation. Genome annotation software is developed with model organisms and does not typically include benchmarking to comprehensively evaluate the quality and accuracy of the final predictions. Plant genomes are particularly challenging with their large genome sizes, abundant transposable elements (TEs), and variable ploidies. This study investigates the impact of genome quality, complexity, sequence read input, and approach on protein-coding gene prediction.

**Methods:** The impact of repeat masking, long-read, and short-read inputs, *de novo*, and genome-guided protein evidence was examined in the context of the popular BRAKER and MAKER workflows for five plant genomes. Annotations were benchmarked for structural traits and sequence similarity.

**Results:** Benchmarks that reflect gene structures, reciprocal similarity search alignments, and mono-exonic/multi-exonic gene counts provide a more complete view of annotation accuracy. Transcripts derived from RNA-read alignments alone are not sufficient for genome annotation. Gene prediction workflows that combine evidence-based and *ab initio* approaches are recommended, and a combination of short and long-reads can improve genome annotation. Adding protein evidence from *de novo assemblies*, genome-guided transcriptome assemblies, or full-length proteins from OrthoDB generates more putative false positives as implemented in the current workflows. Post-processing with functional and structural filters is highly recommended.

**Discussion:** While annotation of non-model plant genomes remains complex, this study provides recommendations for inputs and methodological approaches. We discuss a set of best practices to generate an optimal plant genome annotation, and present a more robust set of metrics to evaluate the resulting predictions.

## INTRODUCTION

The first published plant genome, *Arabidopsis thaliana*, was released in 2000 (Arabidopsis Genome Initiative, 2000). Its small genome size (135Mb) and minimal repeat content stand in stark contrast to the plant species sequenced and assembled today (Kress et al., 2022). NCBI’s (https://www.ncbi.nlm.nih.gov/) genome repository contains genomes of over 900 land plant species, and roughly half of these are assembled to chromosome scale. The total number of complete reference plant genomes has more than doubled in the last five years (Marks et al., 2021). Initiatives like the Open Green Genomes (OGG) (https://phytozome-next.jgi.doe.gov/ogg/), 10KP (Cheng et al., 2018), and the Earth BioGenome Project (Lewin et al., 2022) are improving the phylogenetic representation of plant genomes by sampling underrepresented clades. The plant genomes published today are more likely to be polyploids and/or larger genomes with substantial transposable element content (Sun et al., 2022). The combination of high throughput sequencing advancements, particularly long reads and chromosome conformation capture approaches have enabled the completion of these more challenging assemblies (Pucker et al., 2022).

While genome assembly has seen substantial improvements in accuracy and contiguity, structural annotation remains challenging. This process delineates the physical positions of genomic features, including protein-coding genes, promoters, and regulatory elements. It can be followed by functional annotation, which assigns biological descriptors to the identified features. The accurate classification of these features provides the basis for questions focused on species evolution, population dynamics, and functional genomics. Errors in genome annotation are frequent, even among well-studied models, and are propagated through downstream analyses (Deutekom et al., 2019; Meyer et al., 2020; Salzberg, 2019). In most eukaryotes, genome annotation is challenged by partial conservation of sequence patterns, variable lengths of introns, variable distances between genes, alternative splicing, and higher densities of TEs and pseudogenes (Kersey, 2019; Salzberg, 2019). As a result of these complexities, the structural annotation process requires more advanced informatic tools and skills that support the integration and manipulation of large datasets (Mudge & Harrow, 2016).

Structural and functional genome annotation proceeds in three stages: identifying and masking noncoding regions (repeats); predicting physical positions of gene structures; and assigning biological information to the predictions (Jung et al., 2020). Repeat regions are soft-masked (eg., RepeatMasker (Smit, AFA, Hubley, R & Green, P., 2013-2015) and RepeatModeler2 (Flynn et al., 2020)), which means these regions are indicated but not obscured to annotation software. This is followed by gene prediction, which may be *ab initio* (evidence-free) or evidenced-based. Evidence-based approaches use RNA-Seq and protein sequence similarity search alignments. Evidence-based approaches are often used in combination with *ab initio* (e.g. AUGUSTUS; (Stanke & Waack, 2003)) to generate models that are trained on patterns associated with true genes. Given the advanced state of high throughput transcriptome sequencing, it is common to resolve transcripts from RNA reads through genome-guided approaches, such as StringTie2 (Kovaka et al., 2019). Long-read cDNA sequencing through PacBio and Oxford Nanopore can provide additional resolution and improve the identification of splice variants. When extrinsic evidence from RNA-seq and protein alignments are available, workflow packages like MAKER (Campbell, Holt, et al., 2014; Cantarel et al., 2008; Holt & Yandell, 2011) and BRAKER (Brůna et al., 2021; Hoff et al., 2016, 2019) can assist in training *ab initio* prediction tools. These packages can leverage sequence data from the target species as well as external evidence from closely related species. While these workflows can simplify the integration across external evidence, downstream packages are still required to select or modify the resulting predictions (Banerjee et al., 2021; Gabriel et al., 2021; Haas et al., 2008). Here, we provide a comprehensive evaluation of plant genome annotation workflows, intentionally selecting beyond the typical model species to represent some of the more complex genomes under investigation today. In doing so, we evaluate the impact of repeat-masking using two different implementations of the RepeatModeler2 framework (Flynn et al., 2020). This is followed by exploring the role of read length and accuracy, and the impact of short-read and long-read data. Finally, we examine the contribution of protein evidence, generated from d*e novo* assembly of the RNA inputs and a genome-guided assembly. These variations are examined in the MAKER and BRAKER frameworks to emphasize the importance of defining benchmarks to guide downstream filtering approaches. Finally, the largest and most repetitive genome assessed in this study, *Liriodendron chinense*, was used to demonstrate best practices to refine the predictions.

## METHODS

### Gathering plant genome datasets

Five plant genomes were chosen for this study, including Chinese tuliptree (*Liriodendron chinense*) (Chen et al. 2019), black cottonwood (*Populus trichocarpa* v3) (Tuskan et al. 2006), Chinese rose (*Rosa chinensis*) (Raymond et al. 2018), thale cress (*Arabidopsis thaliana* TAIR 10) (Cheng et al. 2017), and a bryophyte, the common cord-moss (*Funaria hygrometrica*) (Kirbis et al., 2022) (Table S1). The genomes were selected to represent two model systems (*Populus* and *Arabidopsis*) with well curated structural annotations and three non-model systems that exclusively used computational techniques to produce the annotations. Two of these non-models were also more divergent examples, representing the only sequenced member of their genus (*Funaria* and *Liriodendron*). The public assembly and annotation for each species were accessed from NCBI and genome completeness was estimated by searching the genome and annotation for the conserved single-copy orthologs in the Embryophyta odb10 BUSCO v.5.0.0 (Simão et al., 2015). The contiguity of the reference genomes was assessed with Quast v5.0.2 (Gurevich et al., 2013). Published annotation files were summarized with gFACs (Caballero and Wegrzyn, 2019).

Read sets available through NCBI’s Sequence Read Archive (SRA) were accessed to provide transcriptomic evidence for each species and included a variety of tissue types. The Illumina short-read libraries were sequenced with Illumina HiSeq 2500 (100bp paired-end). The read sets included at least four libraries, between 20-82M reads before quality control (QC), and a minimum of 16M reads after QC. Pacific Biosciences Iso-Seq long-reads were accessed for *Populus* and *Liriodendron*, and Oxford PromethION reads were available for *Rosa* and *Arabidopsis*. The read sets for long-read data ranged between 161K-41M total reads per species (Table S2).

### Repeat masking and read alignment

RepeatModeler2 (Flynn et al., 2020) was used to construct repeat libraries with default settings, and repeats were soft masked with the libraries constructed via RepeatMasker v.4.0.6 (Smit et al. 2013-2015). The genomes of *Arabidopsis, Funaria, Populus, and Liriodendron* were additionally masked using RepeatModeler2 with additional LTR identification (-LTRStruct flag). Quality assessment of the Illumina short reads was performed using FastQC v.0.11.7 (Andrews, 2010) before and after trimming low-quality bases. Sickle v.1.33 (Joshi NA, 2011) was used to trim low-quality bases with 50bp as the minimum read length threshold. Single end reads generated post trimming were excluded from RNA alignments and assembly. The trimmed short reads were aligned against their reference genomes using HISAT2 v2.2.0 (Kim et al., 2019). HISAT2 was selected for its performance in recent benchmarking studies and as the aligner of choice for input to Stringtie2 (Corchete et al., 2020; Musich et al., 2021). Long-read RNA data were obtained for four species: *Arabidopsis* and *Rosa* were sequenced with Oxford Nanopore, and *Populus* and *Liriodendron* were sequenced with PacBio Sequel. The long-read data sets were aligned against their respective genomes using Minimap2 v2.1.7 (Li, 2018, 2021).

### Generation of protein evidence

To generate protein evidence, Illumina short reads were assembled *de novo* using Trinity v.2.8.5 with a minimum contig length of 300 bp (Grabherr et al., 2011). The assembled transcriptomes for the multiple libraries were combined, and putative coding regions were predicted using TransDecoder v.5.3.0 (http://transdecoder.github.io). TransDecoder is one of several frame-selection methods available and performs in a comparable manner but not always superior in all metrics (Bolger et al., 2018). For this study, it was selected as the most widely used package for this purpose. Redundancy in the predicted coding regions was reduced after clustering at 98% identity using UCLUST, a clustering algorithm of USEARCH v.9.0.2132 (Edgar, 2010). Frame-selected transcripts shorter than 300 bp were removed. The remaining transcripts were aligned to the genome using GMAP v.2019-06-10 (Wu & Watanabe, 2005). The predicted proteins (from the same Transdecoder run) were aligned to the reference genome using GenomeThreader v 1.7.1 (Gremme, 2014).

To provide protein evidence from genome-guided sources, the previously aligned Illumina short-reads (via HISAT2) were constructed into transcripts with StringTie2 v2.2.0 (Kovaka et al., 2019; M. Pertea et al., 2015). Long-reads were treated similarly, along with a combination of short and long-reads. The predicted transcripts were extracted using gffRead (G. Pertea & Pertea, 2020) and frame selected with TransDecoder. The transcriptome alignment annotation file (gff3) was passed to gFACs for evaluation of gene model statistics. Completeness of the aligned transcripts and protein sequences were estimated using BUSCO. Finally, to provide evidence from external sources (not derived from any transcriptomic inputs), full-length protein evidence from OrthoDB (odb10_plants) was provided for BRAKER/TSEBRA runs.

### Genome annotations

Each genome was tested in four primary open-source annotation softwares to predict gene models (Table 1). Several different runs of BRAKER v.2.1.5 (Hoff et al., 2019) and BRAKER/TSEBRA (Gabriel et al., 2021) were used with various combinations of RNA-Seq (long and short-read inputs) and protein evidence. MAKER v.3.1.3 (Cantarel et al., 2008) was run once with transcript and protein evidence. Finally, StringTie2 (Kovaka et al., 2019), with TransDecoder, was used to generate genome-guided predictions from RNA evidence alone (Table S12).

**Table 1:** Notations for the different runs performed for benchmarking.

| Run |  | *Arabidopsis* | *Funaria* | *Populus* | *Liriodendron* | *Rosa* |
| --- | --- | --- | --- | --- | --- | --- |
| **StringTie2** |  |  |  |  |  |  |
| ST2 (SR) | Short-reads | X | X | X | X | X |
| ST2 (LR) | Long-reads | X |  | X | X | X |
| ST2(SR/LR) | Short and long-reads | X |  | X | X | X |
| **BRAKER** |  |  |  |  |  |  |
| BR (SR) | Short-reads | X | X | X | X | X |
| BR (LR) | Long-reads | X |  | X | X | X |
| BR (SR/LR) | Short and long-reads | X |  | X | X | X |
| BR (SR/RM2+) | Short-reads with additional masking for LTRs | X | X | X | X |  |
| **TSEBRA** |  |  |  |  |  |  |
| TSB (SR/TRINITY) | Short-reads and *de novo* proteins | X | X | X | X | X |
| TSB (SR/ST2) | Short-reads and genome-guided proteins | X | X | X | X | X |
| TSB (LR/ST2) | Long-reads and genome-guided proteins | X |  | X | X | X |
| TSB (SR/LR/ST2) | Short and long-reads and genome-guided proteins | X |  | X | X | X |
| TSB (SR/ST2/RM2+) | Short-reads and genome-guided proteins with additional masking for LTRs | X | X | X | X |  |
| **TSB (SR/OrthoDB)** | **Short-reads and OrthoDB proteins** | **X** | **X** | **X** | **X** | **X** |
| **MAKER** |  |  |  |  |  |  |
| MK (RM2+) | Short-reads with additional masking for LTRs | X | X | X |  |  |

SR- Short reads, LR- Long reads, and RM2+- RepeatModeler2 with the additional repeat masking.

### MAKER annotation

MAKER (MK) was run on the soft-masked reference genomes of *Arabidopsis, Populus*, and *Funaria* with repeats estimated using the additional LTR detection method in RepeatModeler2 (LTRStruct flag; RM2+). This was intended to emulate the MAKER-P (Campbell, Law, et al., 2014) method since the original repeat and pseudogene identification protocols are deprecated. MK (RM2+) was executed (i.e., trained) twice. The annotations derived from MK (RM2+) used protein evidence generated from *de novo* assembled RNA-reads from Trinity. These models were used to train *ab initio* gene prediction software AUGUSTUS v.3.3.3 (Stanke & Waack, 2003) and SNAP v. 2006-07-28 ((Korf, 2004). The Hidden Markov Models (HMMs) trained using AUGUSTUS and SNAP were used along with initial aligned evidence (est2genome and protein2genome parameters) for the second MK (RM2+) run to generate the final gene models.

### Assessment of gene predictions

The quality of genome annotation among different gene prediction methods was evaluated with three primary metrics: (1) the mono-exonic (single-exon) and multi-exonic (multiple exon) ratio; (2) conserved single-copy orthologs queried from the predicted gene models using BUSCO (embryophyta database v10), and (3) gene prediction assessment with EnTAP v0.10.8 (Hart et al., 2020) using a 70% reciprocal functional annotation approach with NCBI’s Refseq Plant and Uniprot databases. The mono:multi ratio was calculated from the gFACs summary report run with default parameters (Caballero & Wegrzyn, 2019). We regard a mono: multi ratio near 0.2 to be ideal and have further validated this with a larger set of model plant genomes (Table S3) (Jain et al. 2008). The gene prediction assessment was recorded as a percentage of sequence similarity hits to the total number of genes. The value of the reciprocal BLAST search will be dependent on the phylogenetic relationships of the target species. While higher annotation rates indicate better gene prediction assessment, species-specific genes will always be missed (Armisén et al., 2008), therefore a minimum annotation rate of 80% is a reasonable threshold to expect when dealing with most non-models. Similarly, a higher BUSCO score indicates a better annotation since BUSCO utilizes OrthoDB to form its conserved sets and the recommended target score is >95% for land plants (Manni et al., 2021; Simão et al., 2015).

The sensitivity and precision of the runs for *Arabidopsis* and *Populus* were assessed using Mikado v2.3.2 (Venturini et al., 2018), by comparing the predicted gene models to the current reference annotations.

### Post-processing filtering

The predicted gene models for *Liriodendron* were taken a step further to refine the genome annotation. Post-process filtering was performed using gFACs and assessed for improvement using BUSCO completeness scores and annotation rates. The mono-exonic and multi-exonic genes predicted for *Liriodendron* were filtered for unique genes. The mono-exonic genes were further filtered for the presence of protein domains using InterProScanv.5.35-74.0 and Pfam (Jones et al., 2014; Quevillon et al., 2005). Multi-exonic genes that did not have an EggNOG or a sequence similarity hit were removed, and the final annotation was assessed using gFACs and EnTAP.

## RESULTS

### Genome sizes, repeats, and published annotations

The genome sizes of the five species assessed represented a 10-fold difference between the smallest genome of *Arabidopsis* (~119 Mb) and the largest of *Liriodendron* (~1.7 Gb) (Fig 1A; Table 2). *Liriodendron* (73.18%) and *Rosa* (60.58%) have higher levels of repeat content, and *Arabidopsis* has the lowest (23.9%). *Arabidopsis* is the most complete chromosome-scale genome, with seven contigs reflecting its five chromosomes and two organellar chromosomes. The other genomes are assembled into pseudochromosomes (except for *Liriodendron*). Once the genomes were downloaded, contigs < 500 bp were removed. The published genome assemblies and annotations were compared in terms of completeness via BUSCO (Fig 1, Table 2). When BUSCO is run in genome mode, it searches the genome for the set of 1614 single-copy orthologs in the embryophyte database. Aside from *Funaria*, which had the lowest completeness score of 82.4%, the remaining plant genomes ranged from 94% to 99%. When we evaluated the published annotations for the same species, and ran BUSCO in protein mode, a slight decrease in completeness was observed in every species except *Funaria* and *Arabidopsis* (Fig 1B). The largest reduction in BUSCO score was observed in *Liriodendron* (98.6% to 75.1%). The discrepancy between the estimated completeness at the genome-level and most of the published annotations speaks to the challenges of achieving an accurate structural annotation.

**Figure 1:**
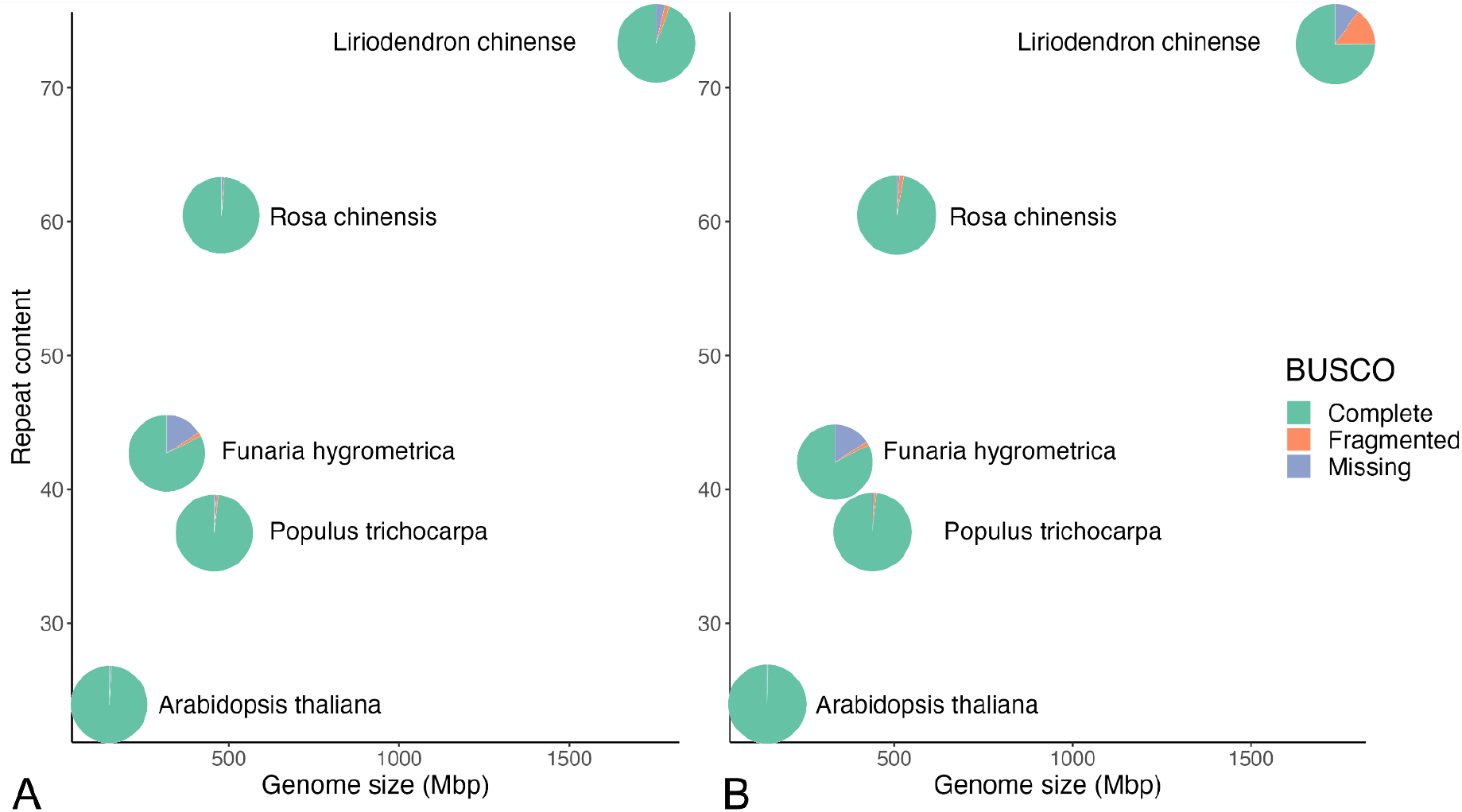
*Genome size, repeat content, and BUSCO completeness for the five plant genomes: Arabidopsis, Populus, Funaria, Rosa, and Liriodendron. Each pie represents the BUSCO completeness. Green denotes the completeness score, orange indicates the fragmented score, and blue indicates the missing score from BUSCO*. (A) *BUSCO scores estimated from the **published assemblies.** (B) BUSCO scores estimated from protein-coding gene predictions from the **published annotations***.

**Table 2:** Genome assembly and annotation statistics for the five published plant genomes

| Species | Genome size | Total scaffolds (chromosomes) | N50 | Repeat content | BUSCO Completeness |  |
| --- | --- | --- | --- | --- | --- | --- |
|  |  |  |  |  | Genome | Annotation |
| *Arabidopsis thaliana* | 119 Mb | 7 (5) | 23.46 Mb | 23.6% | 99.30% | 99.60% |
| *Funaria hygrometrica* | 327 Mb | 687 (26) | 1.48 Mb | 42.35% | 85.60% | 86.60% |
| *Liriodendron chinense* | 1,742 Mb | 3,711 (21) | 3.53 Mb | 73.18% | 98.60% | 75.10% |
| *Populus trichocarpa* | 434 Mb | 1,446 (19) | 19.47 Mb | 35.90% | 98.80% | 98.30% |
| *Rosa chinensis* | 515 Mb | 55 (7) | 69.64 Mb | 60.53% | 98.80% | 97.30% |

RepeatModeler2 (RM2) with and without the LTRStruct package (the additional LTR masking module) (Flynn et al. 2020) was used to soft mask repeats in four of the genomes. The increase in repeat content was marginal in all species, ranging from 1% in *Funaria* to 5% in *Populus*. Comparisons using the LTRStruct flag were denoted as RM2+ (Table S4).

### Transcriptome evidence

For the subsequent genome annotation analysis, the Illumina RNA short reads were first aligned to the genome. All libraries, ranging from four to 20 per species, aligned at over 97%, apart from *Rosa* (92%) (Table 3; Table S5). Long-read RNA libraries were aligned with Minimap2 for four species: *Arabidopsis* (Nanopore reads at 97.1%), *Populus* (Iso-Seq reads at 92.01%)*, Liriodendron* (Iso-Seq reads at 95.5%), and *Rosa* (Nanopore reads at 99%). The N50s for the long reads range from 976 bp in *Rosa* to 4.6 Kb in *Liriodendron* (Table S6).

**Table 3:**
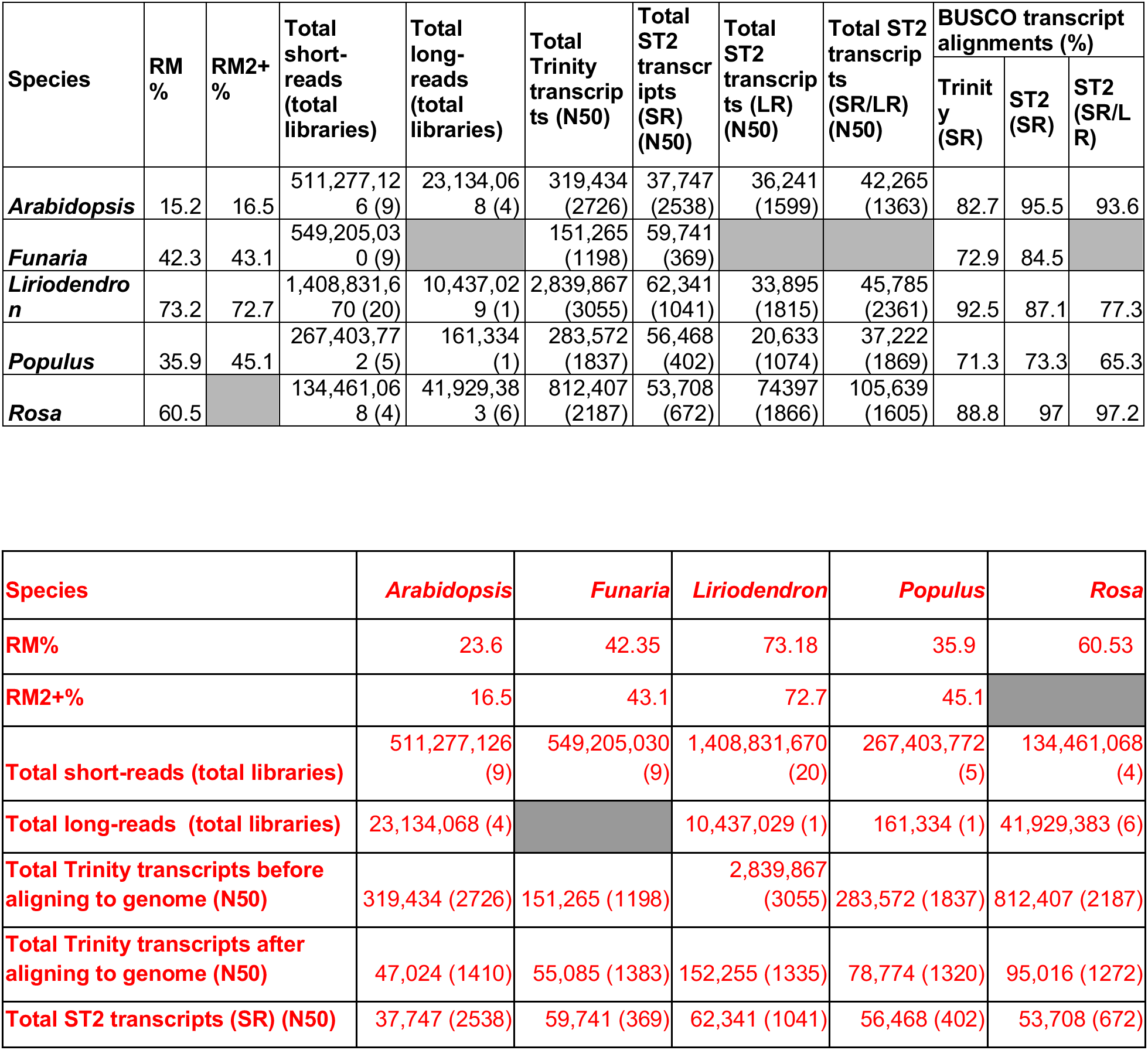
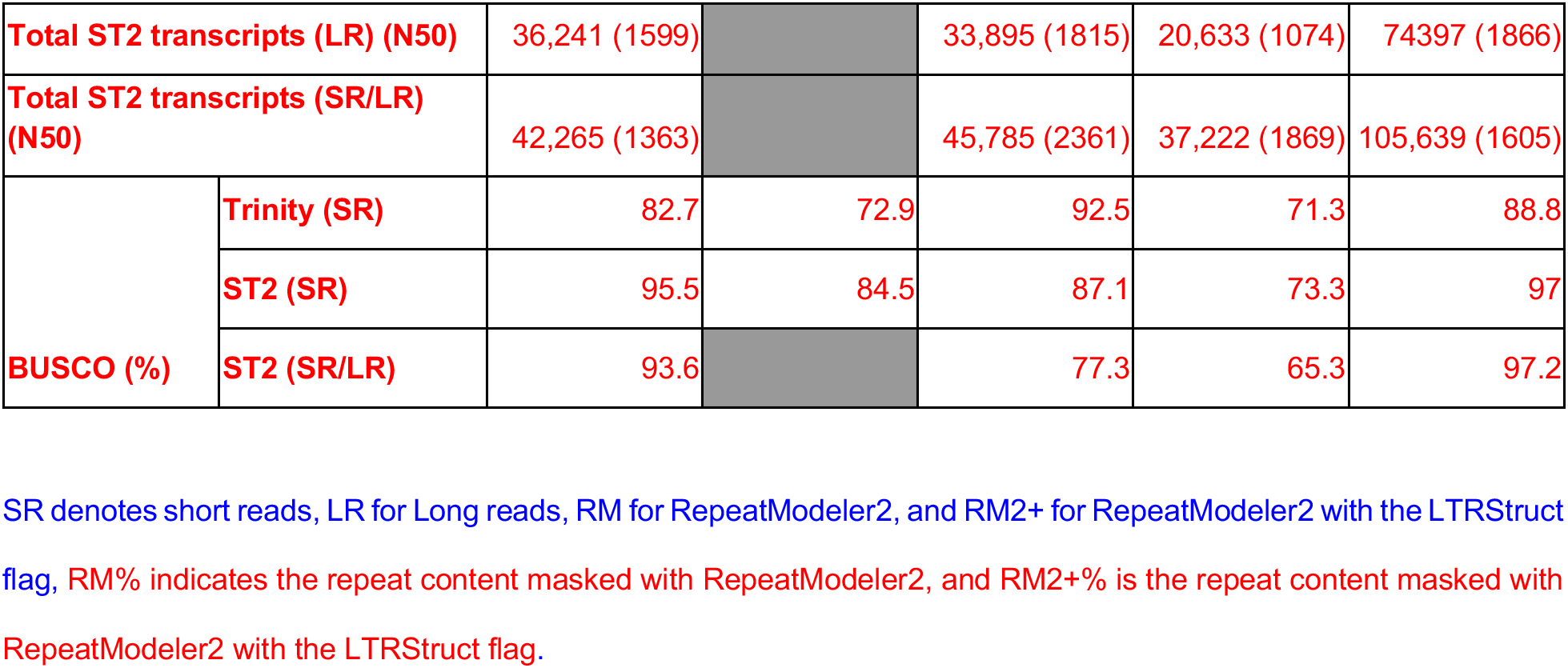
Comparison between genome-guided (StringTie2-ST2) and de novo (Trinity) genome annotations.

| Species | RM % | RM2+ % | Total short-reads (total libraries) | Total long-reads (total libraries) | Total Trinity transcripts (N50) | Total ST2 transcripts (SR) (N50) | Total ST2 transcripts (LR) (N50) | Total ST2 transcripts (SR/LR) (N50) | BUSCO transcript alignments (%) |  |  |
| --- | --- | --- | --- | --- | --- | --- | --- | --- | --- | --- | --- |
|  |  |  |  |  |  |  |  |  | Trinity (SR) | ST2 (SR) | ST2 (SR/LR) |
| *Arabidopsis* | 15.2 | 16.5 | 511,277,126 (9) | 23,134,068 (4) | 319,434 (2726) | 37,747 (2538) | 36,241 (1599) | 42,265 (1363) | 82.7 | 95.5 | 93.6 |
| *Funaria* | 42.3 | 43.1 | 549,205,030 (9) |  | 151,265 (1198) | 59,741 (369) |  |  | 72.9 | 84.5 |  |
| *Liriodendron* | 73.2 | 72.7 | 1,408,831,670 (20) | 10,437,029 (1) | 2,839,867 (3055) | 62,341 (1041) | 33,895 (1815) | 45,785 (2361) | 92.5 | 87.1 | 77.3 |
| *Populus* | 35.9 | 45.1 | 267,403,772 (5) | 161,334 (1) | 283,572 (1837) | 56,468 (402) | 20,633 (1074) | 37,222 (1869) | 71.3 | 73.3 | 65.3 |
| *Rosa* | 60.5 |  | 134,461,068 (4) | 41,929,383 (6) | 812,407 (2187) | 53,708 (672) | 74,397 (1866) | 105,639 (1605) | 88.8 | 97 | 97.2 |

| Species | *Arabidopsis* | *Funaria* | *Liriodendron* | *Populus* | *Rosa* |
| --- | --- | --- | --- | --- | --- |
| RM% | 23.6 | 42.35 | 73.18 | 35.9 | 60.53 |
| RM2+% | 16.5 | 43.1 | 72.7 | 45.1 |  |
| Total short-reads (total libraries) | 511,277,126 (9) | 549,205,030 (9) | 1,408,831,670 (20) | 267,403,772 (5) | 134,461,068 (4) |
| Total long-reads (total libraries) | 23,134,068 (4) |  | 10,437,029 (1) | 161,334 (1) | 41,929,383 (6) |
| Total Trinity transcripts before aligning to genome (N50) | 319,434 (2726) | 151,265 (1198) | 2,839,867 (3055) | 283,572 (1837) | 812,407 (2187) |
| Total Trinity transcripts after aligning to genome (N50) | 47,024 (1410) | 55,085 (1383) | 152,255 (1335) | 78,774 (1320) | 95,016 (1272) |
| Total ST2 transcripts (SR) (N50) | 37,747 (2538) | 59,741 (369) | 62,341 (1041) | 56,468 (402) | 53,708 (672) |

|  |  |  |  |  |  |  |
| --- | --- | --- | --- | --- | --- | --- |
| **Total ST2 transcripts (LR) (N50)** |  | 36,241 (1599) |  | 33,895 (1815) | 20,633 (1074) | 74397 (1866) |
| **Total ST2 transcripts (SR/LR) (N50)** |  | 42,265 (1363) |  | 45,785 (2361) | 37,222 (1869) | 105,639 (1605) |
| **BUSCO (%)** | **Trinity (SR)** | 82.7 | 72.9 | 92.5 | 71.3 | 88.8 |
|  | **ST2 (SR)** | 95.5 | 84.5 | 87.1 | 73.3 | 97 |
|  | **ST2 (SR/LR)** | 93.6 |  | 77.3 | 65.3 | 97.2 |

SR denotes short reads, LR for Long reads, RM for RepeatModeler2, and RM2+ for RepeatModeler2 with the LTRStruct flag, RM% indicates the repeat content masked with RepeatModeler2, and RM2+% is the repeat content masked with RepeatModeler2 with the LTRStruct flag.

### Transcript-derived annotations

The reads were assembled using StringTie2 (ST2) and Trinity. Trinity *de novo* assemblies of the Illumina short reads generated longer transcripts, with N50s ranging from 1.2 Kb (151,265 transcripts in total) in *Funaria*, to 3.06 Kb (2,839,867 transcripts) in *Liriodendron*. Among genome-guided assemblies with StringTie2 (ST2(SR)), the range was much smaller, with N50s ranging from 369 bp (59,741 transcripts in total) in *Funaria* to 2.54 Kb (37,747 transcripts) in *Arabidopsis* (Table S6). The StringTie2 (ST2(LR) and ST2(SR/LR)) range was longer, with N50s ranging from 1.07 Kb (20,633 transcripts in total) in *Rosa* ST2(LR) to 2.36 Kb (45,785 transcripts) in *Liriodendron* ST2 (SR/LR) (Table 3; Table S6). The StringTie2 and Trinity transcripts were aligned back to the genome using GMAP after frame-selection. BUSCO scores for the aligned transcriptomes derived from short read data, run in transcriptome mode, ranged from 73% in *Funaria* to 83% in *Rosa* for Trinity, and 73% in *Liriodendron* to 97% in *Rosa* using StringTie2 (Table 3). The BUSCO scores were the lowest for the ST2 (LR) runs across all species as compared to the other StringTie2 only runs. For the ST2 (SR/LR), the BUSCO scores were lower than ST2 (SR), except for *Rosa*, where the ST2 (SR/LR) was 97.2% as opposed to 97% in ST2 (SR). In all species, ST2 (SR/LR) had higher BUSCO scores than ST2 (LR). Despite Trinity producing more than double the total transcripts than StringTie2, the BUSCO completeness score of most StringTie2 runs were much higher than that of Trinity. *Liriodendron* remained the only exception with a slightly higher BUSCO score from Trinity.

*Arabidopsis* and *Populus* were further evaluated with Mikado to compare the sensitivity and specificity of published annotations (Fig 4B, Table S7). Overall, StringTie2 predictions had higher sensitivity and precision rates compared to the Trinity runs. From this point, Trinity was excluded, and StringTie2 runs were compared against BRAKER and TSEBRA predictions (Fig 4B).

The mono: multi ratios produced by StringTie2 ranged from 0.15 in *Populus* (ST2 (LR)) to 0.53 in *Liriodendron* (ST2 (LR)), which were an improvement over the mono: multi ratios produced from the BRAKER annotations that ranged from 0.37 in *Arabidopsis* (BR (LR)) to 1.27 in *Funaria* (BR (SR/RM2+)). The BUSCO scores of the proteins predicted from BRAKER were generally higher than the BUSCO scores from StringTie2. For example, *Arabidopsis* StringTie2 runs range from 85% (ST2 (LR)) to 95.5% in ST2 (SR), and BRAKER runs ranged from 94% (BR (LR)) to 95.9% (BR (SR)). However, some runs are comparable, the ST2 (SR) run with a BUSCO score of 95% was like the BR (SR) run at 95% and the BR (SR/RM2+) run at 95% in *Arabidopsis*. StringTie2 predicted models had a higher annotation rate, in general, compared to BRAKER. For example, the EnTAP annotation rate in *Funaria* was just over 40% post BRAKER but was near 60% from the StringTie2 runs (Fig 2).

**Figure 2:**
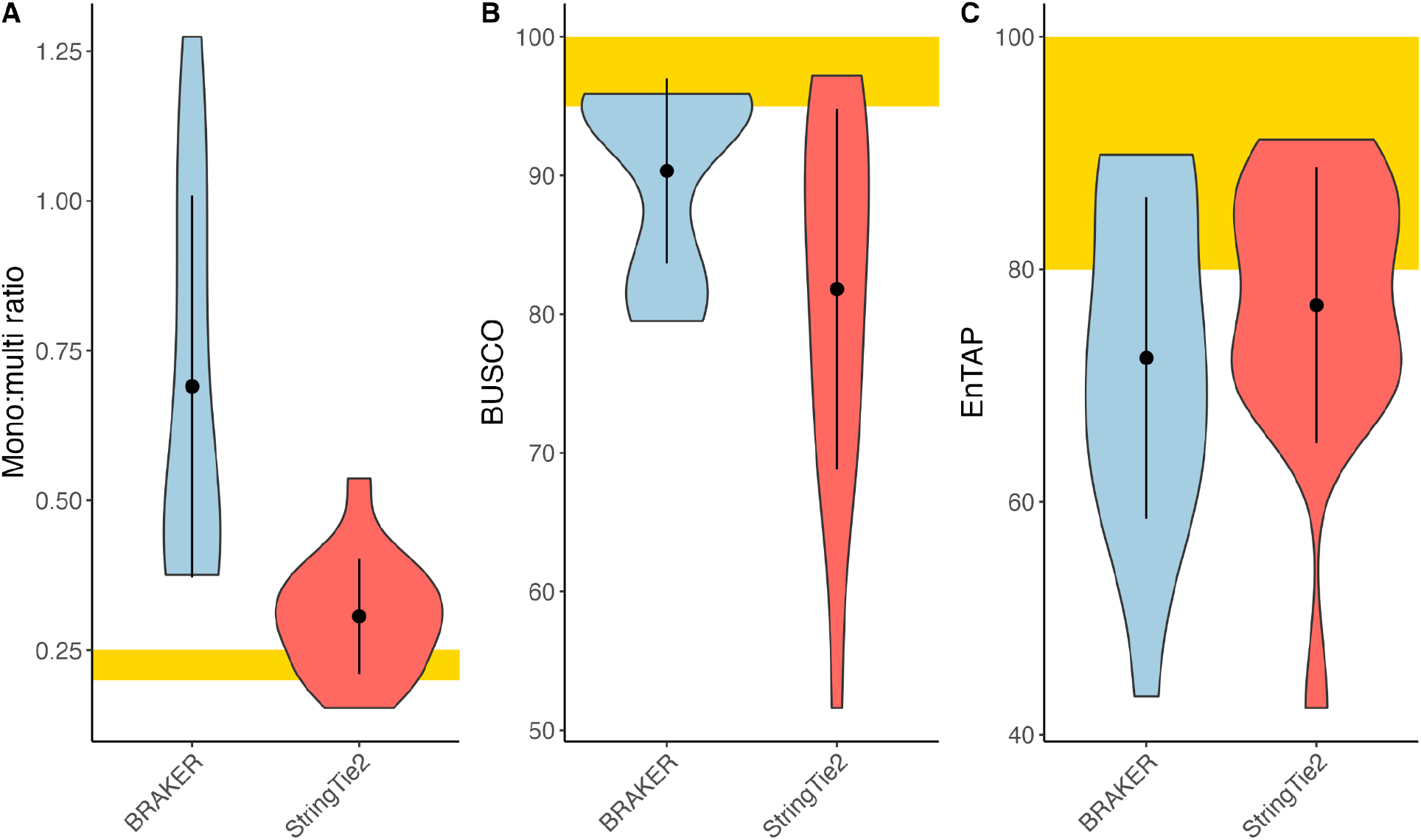
Comparing metrics between BRAKER (blue) and StringTie2 (red) predictions. (A) mono:multi ratios, (B) BUSCO comparisons, and (C) EnTAP annotation rates of the gene models. The yellow region indicates the ideal value for each of the metrics.

### Genome annotation with MAKER

To replicate MAKER-P’s repeat pipeline, the RM2+ genome was used for *Arabidopsis, Populus*, and *Funaria* for the MAKER runs. BUSCO completeness was low, compared with BRAKER runs, and ranged from 19.6% in *Populus* to 90.4% in *Arabidopsis* (Fig 3A). The mono: multi ratio of MAKER(RM2+) for *Arabidopsis* was comparable to the BRAKER runs for the same species (0.22 for BR (SR) and BR (SR/RM2+), 0.24 for BR (LR), and 0.23 for BR (SR/LR)). The MK (RM2+) predictions for the total number of genes in *Arabidopsis* and *Funaria* were in the expected range for these species, 22K and 44K genes, respectively, whereas only 7K genes were predicted for *Populus*. The gene lengths ranged from 1.8 Kb in *Funaria* to 2.3 Kb in *Arabidopsis* (Table S8). The best run for MK (RM2+) was for *Arabidopsis*, with a mono: multi ratio of 0.22 and a BUSCO score of 90.4%. On the other hand, the mono: multi ratio for *Populus* was 0.07, and the BUSCO score was 19.6%.

**Figure 3:**
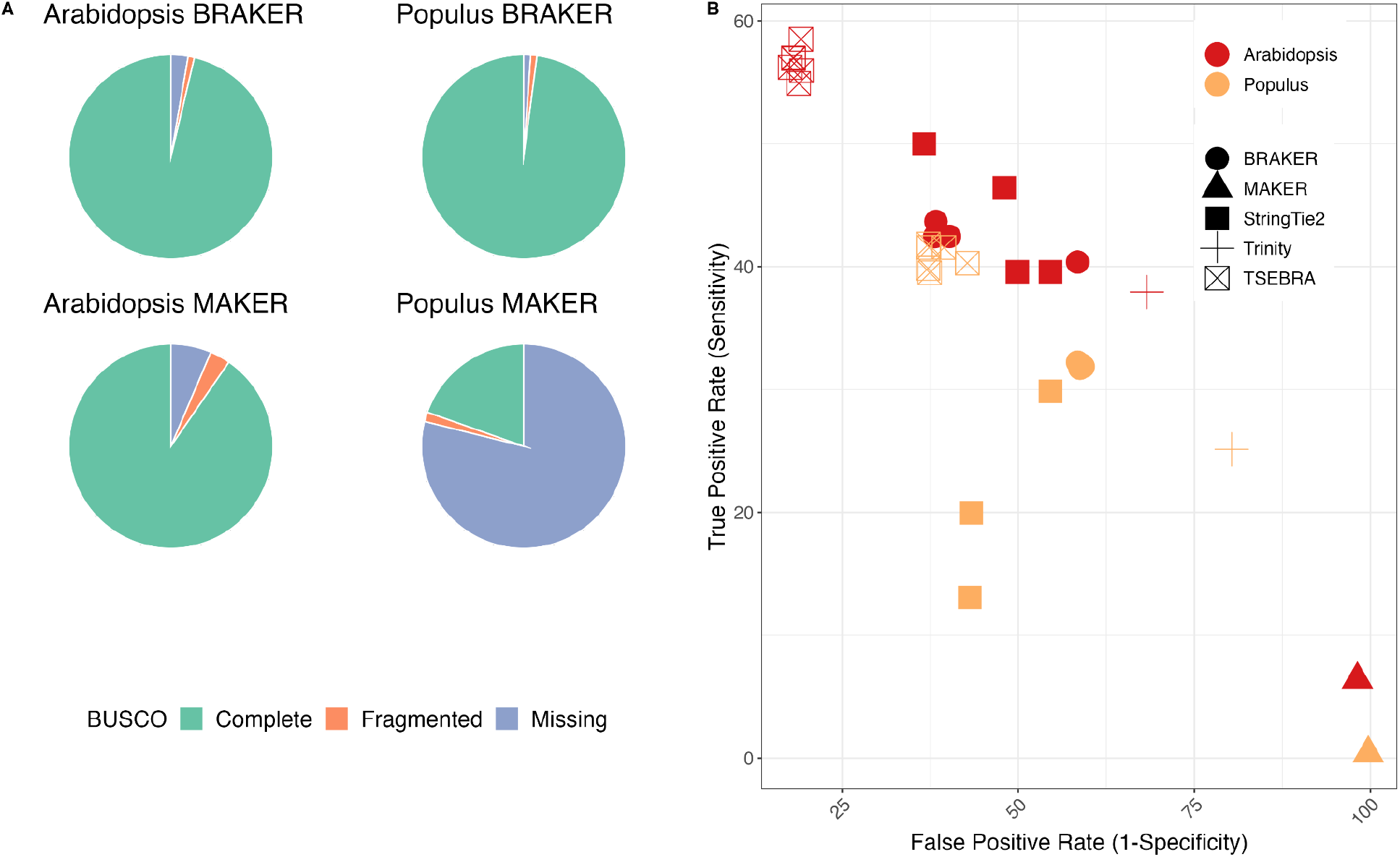
*Comparison of BUSCO, sensitivity, and false positive rates between the Arabidopsis and Populus annotations (Table S10). (A) BUSCO completeness scores for the MK(SR/RM2+) and BR(SR/RM2+) runs of Arabidopsis and Populus, green denotes the completeness score, orange indicates the fragmented score, and blue indicates the missing score from BUSCO (B) False positive rates and sensitivity scores from Mikado against published annotations for Arabidopsis (red color) and Populus (gold color) for the MAKER, BRAKER, TSEBRA, Trinity, and StringTie2 runs. The scores were assessed using MIKADO*. Multiple points per run reflect differences in input read type and repeat-masking.

**Figure 4:**
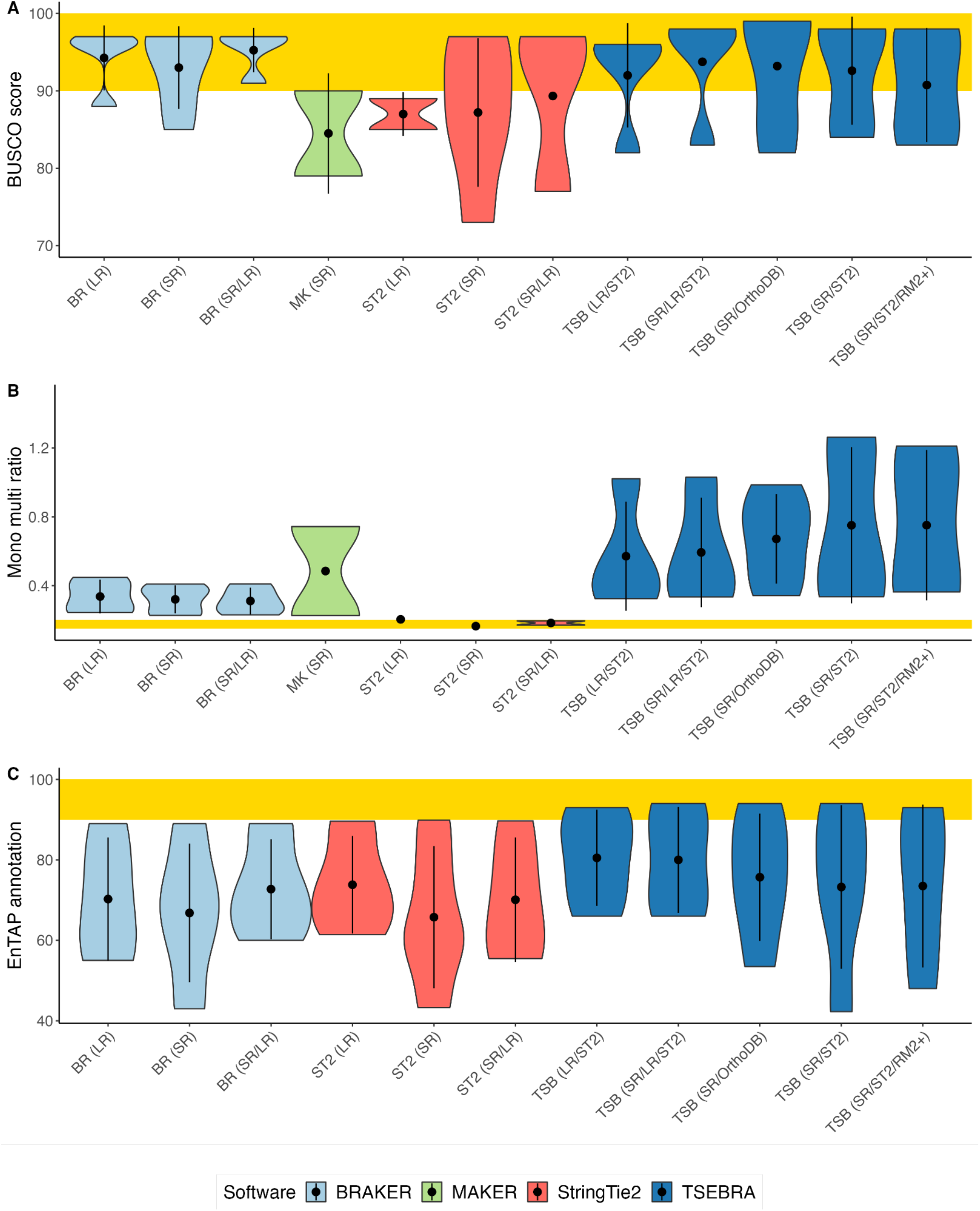
Comparison of BUSCO completeness scores (A), mono:multi ratios (B), and EnTAP annotation rates (C) across all species between the runs of different input types and software, i.e., MAKER (MK-green) BRAKER (BK-light blue), TSEBRA (TSB-dark blue) and StringTie2 (ST2-red). The yellow rectangle represents the target scores for each benchmark. RM2+-RepeatModeler2 with LTRStruct.

The model systems, *Arabidopsis* and *Populus*, further were evaluated with Mikado to compare the sensitivity and specificity of the published annotations (Fig 3B; Table S7). The sensitivity and precision scores for gene predictions were the lowest from MAKER, followed by Trinity, and were the highest from TSEBRA. StringTie2 and BRAKER yielded similar sensitivity and specificity scores for *Arabidopsis*, whereas for *Populus* the sensitivity score was lower than those from BRAKER runs. Given its overall low scores, MAKER was excluded from the subsequent comparisons. It should be noted, however, that protocols are available that recommend using GeneMark in MAKER. With this addition to the training protocol, MAKER was shown to have a higher accuracy in prediction (Brůna et al., 2021).

### Genome annotation with BRAKER

In general, BUSCO scores were improved in the BRAKER and TSEBRA runs, with the TSB(SR/OrthoDB) predictions scoring the highest. The best BUSCO scores were from TSB(SR/OrthoDB), ranging from 98.9% (TSB; SR/OrthoDB) in *Arabidopsis* to 82.2% in *Funaria*. Overall, the TSEBRA runs fared better with the annotation rates ranging from 99% in *Arabidopsis* (TSB; SR/OrthoDB) to 53% in *Funaria*. On the other hand, the TSEBRA runs had the worst mono:multi ratios. For example, TSB (SR/ST2/RM2+) has the highest ratio at 1.27 for *Funaria* (Fig 4).

Overall, the gene models generated by BRAKER for *Arabidopsis* performed similarly across runs when evaluated by BUSCO scores. The mono: multi ratios across BRAKER runs ranged between 0.23 to 0.39, and the annotation rates were consistently above 95%.

The annotation rates for *Funaria* were lower than expected, 43% for BR (SR) and 53% for nearly all methods that included protein-evidence (TSB (SR/TRINITY), TSB (SR/OrthoDB)). The BUSCO completeness scores of about 85% post BRAKER are comparable to those from StringTie2.

In the case of *Liriodendron’s* BRAKER and TSEBRA runs, the mono:multi ratios were more variable when compared to the StringTie2 runs, which ranged from 0.34 BR(SR) to 1.04 with BR (SR/RM2+). The annotation rates for each run were around 75%, with BUSCO scores from 83% with TSB (SR/LR/ST2) to 90.8% for BR (SR). *Populus* gene models post-BRAKER without protein had mono:multi scores around 0.24, and with TSEBRA, the ratio ranged from 0.4 to 0.5. Annotation rates also differed between BR(SR) (75%) and TSB (SR/ST2) (88%). *Rosa* had overall consistent scores for BUSCO post BRAKER, ranging around 96%. TSEBRA runs had higher mono:multi ratios (0.75) compared to the BRAKER runs (0.37) (Table S8).

### Annotation with long reads

For BRAKER runs, the predicted gene lengths from the long-reads were comparable to those based on short-reads, except for some runs with long-reads within *Populus*. The average gene length for BRAKER with long-reads for *Populus* ranges from 2.7 Kb to 3.4 Kb, although some transcripts exceed 6 Kb. The longest average gene lengths were observed in *Liriodendron* (9.3 Kb-BR; SR/LR). The inclusion of long-reads (exclusively) did not improve BUSCO completeness for any species, apart from *Arabidopsis*, where the BR (LR) BUSCO completeness was 1% higher than the BR (SR) run. The increase in BUSCO completeness in *Arabidopsis* could be due to the large number of long-reads included (23M across four libraries). However, the quality of genome annotation does not seem correlated with depth of long-read sequencing; for example, *Rosa* had more reads (41M across 6 libraries), and the BR (LR) run had a similar BUSCO score to BR (SR) (96%). It should be noted that the long reads for *Arabidopsis* and *Rosa* were sequenced with ONT. The ONT reads had higher mapping rates, compared to Iso-Seq, to their respective genomes, 97.1% in *Arabidopsis* and 99% in *Rosa* (Table S5). The long-read inputs, regardless of depth or type, lowered the BUSCO completeness (up to 10%) across all ST2 (LR) runs (Table S9). Finally, we note that the combination of short-reads and long-reads (BR; SR/LR and TSB; SR/LR/ST2) is comparable to the BR (SR) reads in terms of BUSCO completeness. Exceptions in this case are *Arabidopsis* and *Rosa*, with marginally higher BUSCO completeness scores. BR (SR/LR) runs produced more genes (in total) for *Arabidopsis, Populus*, and *Rosa* in comparison to BR (SR); however, in the TSB (SR/LR/ST2) runs, the total number of genes were lower compared to BR (SR), apart from *Rosa*. The annotation rate in BR (SR/LR) of *Liriodendron* was higher than BR (SR). The combination of proteins with SR and LR (TSB(SR/LR/ST2)) resulted in higher annotation rates across all species. The combination of SR and LR increased mono: multi ratios, therefore, resulted in worse mono: multi ratios across all species.

### Refining the genome annotation for Liriodendron

The BRAKER runs for *Liriodendron* were filtered with gFACs and InterProScan to remove unlikely gene models (Table 4). The number of mono-exonic genes was drastically reduced post-filter with InterProScan. Across all runs, the mono-exonic genes numbered 11K to 25K. After removing mono-exonics without a protein domain annotation from the Pfam database, they decreased from 13K to 5K. The decrease in false positive mono-exonics resulted in an improved mono:multi ratio that range from 0.16 for BR (SR) and BR (SR/RM2+), 0.16 and 0.23 for the StringTie2 runs to 0.43 for the TSEBRA runs. The BUSCO scores decreased slightly post-filtering (1-2%). EnTAP annotation rates ranged from 66% to 86%, with the TSB (SR/OrthoDB), which is an improvement from 59% to 68% pre-filtering.

**Table 4:** Gene model statistics for Liriodendron after two rounds of structural and functional filters.

| *Liriodendron*<br>Annotation | Total genes | Mono:Multi Ratio | BUSCO % | EnTAP % |
| --- | --- | --- | --- | --- |
| Published annotation | 35261 | 0.7 | 75.1 | 63 |
| **Mono-exonic filters** |  |  |  |  |
| BR (LR) | 39031 | 0.21 | 87.4 | 69 |
| BR (SR) * | 41065 | 0.16 | 90.2 | 66 |
| BR (SR/LR) * | 40420 | 0.16 | 90.3 | 67 |
| BR (SR/RM2+) | 40740 | 0.17 | 88.2 | 67 |
| ST2 (SR) | 51804 | 0.16 | 86.5 | 80 |
| ST2 (LR) | 27012 | 0.23 | 65 | 84 |
| ST2 (SR/LR) | 36345 | 0.24 | 70.6 | 82 |
| TSB (LR/ST2) | 33132 | 0.43 | 82.3 | 84 |
| TSB (SR/LR/ST2) | 33964 | 0.43 | 82.4 | 84 |
| TSB (SR/ST) | 32898 | 0.41 | 83.4 | 84 |
| TSB (SR/ST2/RM2+) | 33637 | 0.45 | 82.8 | 84 |
| TSB (SR/TRINITY) | 34646 | 0.42 | 84 | 83 |
| TSB (SR/OrthoDB) | 33667 | 0.43 | 86.5 | 86 |
| **+Multi-exonic filters** |  |  |  |  |
| BR (SR) | 30219 | 0.24 | 90.3 | 81 |
| BR (SR/LR) | 30035 | 0.23 | 86.9 | 87 |

\* denotes the two best annotation sets
BR- BRAKER, ST2- StringTie2, TSB- TSEBRA, SR- Short reads, LR- Long-reads, RM2+- RepeatModeler2 with LTRStruct. BUSCO % shows the completeness percentage, and EnTAP % is the annotation rates after reciprocal BLAST.

In terms of BUSCO completeness and mono:multi ratios, the two best performing runs (BR (SR) and BR (SR/LR)) were further filtered (Table 4). In this step, multi-exonic genes without an EggNOG alignment or a sequence similarity alignment through EnTAP were removed. These filtered models were re-assessed for mono:multi ratio, BUSCO completeness, and EnTAP annotation. The BUSCO completeness remained the same for BR(SR), but not for BR(SR/LR). The EnTAP annotation increased from 66% to 81% in BR (SR), and 67% to 87% in BR (SR/LR).

## DISCUSSION

BRAKER (Hoff et al., 2020) and MAKER (Cantarel et al., 2008) are currently the most popular eukaryotic structural annotation tools, cited 475 and 1,010 times (since 2021, as referenced in Google Scholar). Processes that select from multiple *ab initio* or aligned forms of evidence are gaining popularity as well though they add both time and complexity to the analyses (FINDER cited 22 times, (Banerjee et al., 2021); EVidenceModeler cited 381 times (Haas et al., 2008)). Finally, as high-throughput transcriptomics, in the form of both short and long-read evidence become more accessible, rapid approaches like StringTie2 (cited 451 times (Kovaka et al., 2019)) are occasionally used as the exclusive approach, though more often, used in combination with the options listed above.

Regardless of the methods selected, recently published benchmarks are challenged to achieve high values for gene sensitivity in larger genomes (Brůna et al., 2021). Within smaller and less complex model systems such as *C. elegans* and *D. melanogaster, ab initio* prediction results in gene sensitivity 49.8% and 59.5%, respectively (Brůna et al., 2021). In well studied complex organisms, such as humans, gene level sensitivity and specificity hovers at 48% and 43%, respectively (Banerjee et al., 2021). While generating benchmarks with model systems (*A. thaliana, C. elegans*, and *D. melanogaster*) provides more reliable metrics for comparison, they are infamous for not fully representing the diversity of their respective clades (Chang et al., 2016).

This study focused on four gene prediction workflows: StringTie2, MAKER, BRAKER, and BRAKER/TSEBRA, and examined the process across a variety of evidence inputs. Both model and non-model plant genomes were considered to highlight the challenges and reinforce the need for downstream filtering.

### Genome annotation benchmarks for both models and non-models

Among plant genomes, the total number of genes is relatively conserved and ranges from 20,000 to just over 40,000. As such, total gene number provides an accessible preliminary benchmark. However, the number of genes in the reference annotation fails to assess the overall quality of the annotation. To measure this, we should consider additional metrics. Here, we describe the utility of BUSCO score, mono-exonic: multi-exonic ratio, and sequence similarity assessment.

BUSCO allows us to identify complete, duplicated, fragmented, and missing single-copy orthologs shared by most seed plants (Simão et al. 2015; Seppey, Manni, and Zdobnov 2019). This provides a reliable benchmark in the absence of a high quality reference annotation and poor BUSCO scores are immediately indicative of a larger issue. However, a high BUSCO score is not sufficient to estimate the quality of an annotation (Fig 4B). Six of the 18 BRAKER runs and four of the 17 StringTie2 runs exceeded 95% completeness. However, total gene number, gene length, and structure varied considerably.

Repeat content, especially in the form of LTRs, and pseudogenes can lead to inflated gene model estimates, especially in the form of mono-exonic genes (Scott et al., 2020; Trouern-Trend et al., 2020). We expect that eukaryotes maintain 20% or less of their gene space as mono-exonics (Jain et al., 2008; Table S3). Although the BUSCO scores were consistent, we note tremendous variation in mono- to multi-gene model ratios post-BRAKER. In practice, having a worse mono: multi ratio is preferable to having a lower BUSCO score, since missing genes, especially those thought to be conserved, cannot be easily rectified, while putative false positives can potentially be filtered through other means.

Sequence similarity search metrics are more complex to interpret, but when used with high-quality and curated databases that contain full-length proteins (e.g. NCBI RefSeq), can provide a benchmark. Specifically, a reciprocal BLAST search requires that both the query and target in the search retain a minimum level of coverage in the alignment. For new plant genomes, that are in the darkest corners of the tree of life, this might be a less reliable metric. For species that may fare poorly in database comparisons, searches for protein domains can provide some level of confidence and we demonstrate this as a filter to reduce the mono-exonics in *Liriodendron*.

### Masking repeats in plant genomes: Repeat masking is important but may not require additional LTR resolution to improve performance

Plant genomes typically contain many repeats, mostly in the form of transposable elements (TEs), averaging around 46% (Luo et al., 2022). Given the abundance of TEs in genomes, it is important to mask these in advance of gene prediction. Soft masking involves changing nucleotides identified as repeats to lowercase (Yandell & Ence, 2012), signaling downstream programs to ignore these sequences. Of the five genomes included in this study, *Liriodendron* had the largest genome size and repeat content. Running downstream analyses on an unmasked genome of *Liriodendron* resulted in a 4-fold increase in gene predictions (Fig 5A). Many repeats were identified as putative gene models, resulting in a large increase of total number of genes (Fig 5B).

**Figure 5:**
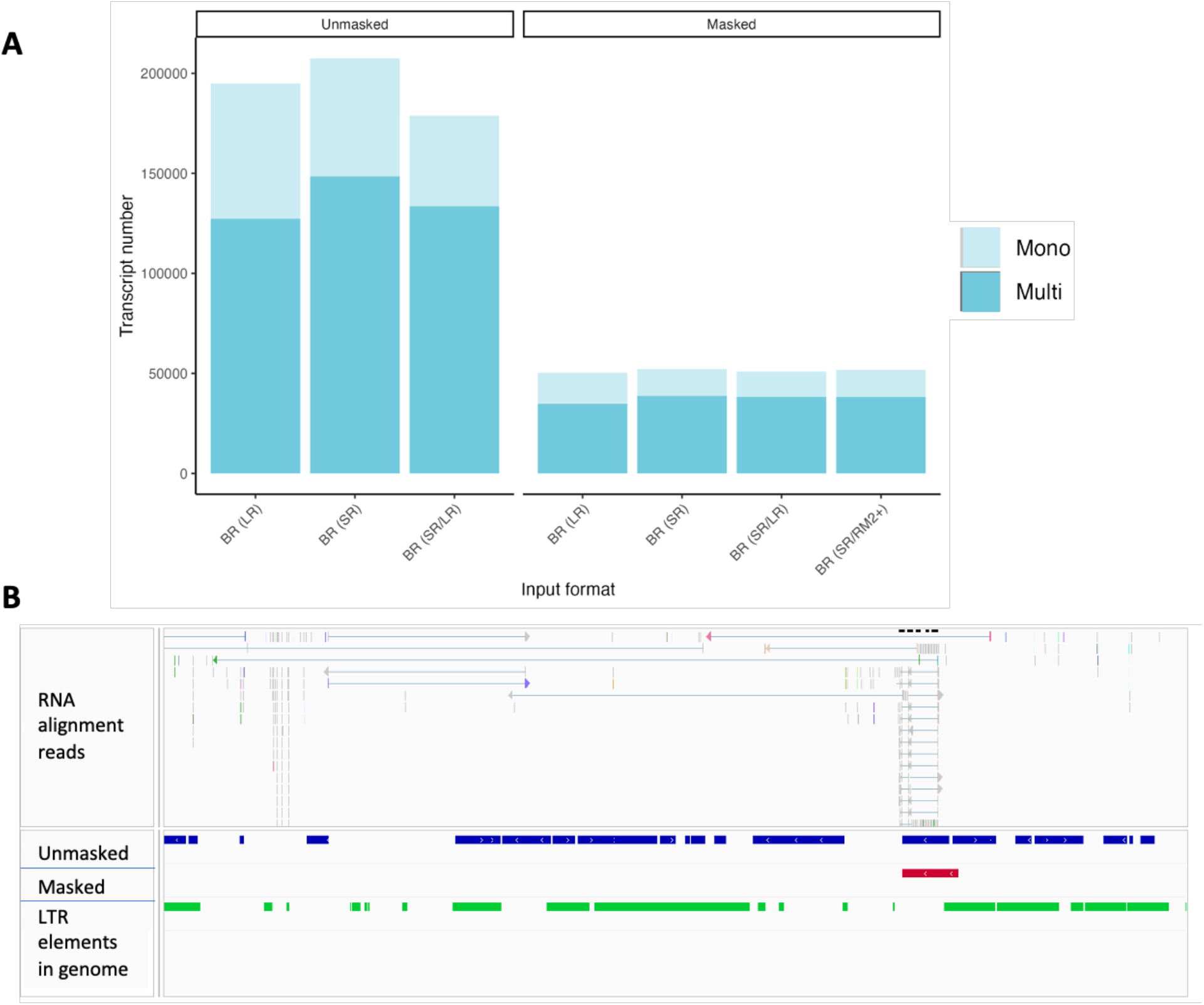
(A) The effect of soft-masking on gene prediction in Liriodendron (Table S13). Performing structural annotation on the unmasked Liriodendron genome results in inflation in the mono and multi-exonic genes. Blue denotes the BRAKER (BR) runs for both genomes, SR denotes short-reads, and LR denotes long reads. The lighter shade represents mono-exonics, and the darker shade represents the multi-exonics. (B) More genes predicted using the unmasked genome (blue), as compared to only one gene predicted in this region with the masked genome (red). The green track shows the LTR elements in the genome as identified by RepeatModeler2. The RNA alignment reads show a read pile-up at the predicted gene (masked track).

RepeatModeler2 is a widely used tool for TE discovery (Flynn et al., 2020). The recent release of RepeatModeler2 includes an optional module for more robust LTR structural detection (LTRStruct module) that includes the LTRharvest (Ellinghaus et al., 2008), LTRDetector (Valencia & Girgis, 2019), and LTR_retriever packages (Ou & Jiang, 2018). This is particularly useful in identifying more divergent LTRs in the genome that may exist in fewer copies (Ou & Jiang, 2018; Valencia & Girgis, 2019). Among the default packages included, RepeatScout serves as a fast method to detect young and abundant repeat families in the genome. RECON, on the other hand, is more computationally intensive and is sensitive enough to detect older TE families. The LTRStruct module is run on the unmasked genome to identify LTR families that may be redundant with the families identified by the default package. This creates redundancy that is resolved through clustering with CD-HIT (Flynn et al., 2020).

In the four species compared, additional repeat masking did not significantly improve gene predictions (Table S9; Fig 4). The mono: multi ratios across species were consistent before and after additional LTR masking (Fig 5A). The BUSCO completeness scores remained relatively the same, with BR (SR/RM2+) being 1% higher than BR (SR) in *Arabidopsis, Funaria* and *Populus*. The marginal improvement observed in these genomes could be related to the structure and type of LTRs, for example, better identification of divergent Ty1-copia elements described in the *Funaria* genome (Kirbis et al., 2022). While we did not include genomes with excessive repeat estimates (>70%), our results indicated that the optional LTRStruct module was not beneficial.

### Genome-guided transcriptome assembly for annotation: Transcripts derived directly from alignments are not sufficient to annotate reference genomes

Transcriptome assemblers are designed to work with primarily short RNA-Seq reads to construct full-length transcripts. In the presence of a high-quality reference genome, genome-guided approaches are preferred as the reads are aligned directly to the target genome in advance. Aligned RNA evidence provides resolution on exon boundaries, and aids in the identification of splice variants. *De novo* approaches build graph models directly from the short (or long) reads to generate transcripts. The latter is much more challenging, computationally intensive, and prone to error.

We compared the accuracy of the annotations produced by StringTie2; *de novo* assembled transcripts with Trinity to annotations produced by BRAKER. The selected packages are top performers when compared in their respective categories of genome-guided and *de novo* transcriptome assembly (Sahraeian et al., 2017; Venturini et al., 2018). As expected, Trinity produced a higher number of transcripts than StringTie2, and BUSCO completeness was consistently lower (Table 3), except for *Liriodendron*. The gene models generated by StringTie2 were more numerous than the BRAKER gene models, more than expected for each species. It should be noted, however, that StringTie2 identifies splice variants by generating a splice graph and resolving conflict between multiple potential splice sites (Kovaka et al., 2019), whereas BRAKER trains an internal algorithm GeneMark-ET to find specific genes with complete support among all introns to be further used in training Augustus (Hoff et al., 2019).

StringTie2 runs resulted in lower BUSCO completeness when compared to BRAKER and/or TSEBRA runs (Fig 4A; Table S11). This outcome is supported by the lack of *ab initio* prediction with genome-guided approaches. Inflated mono-exonic predictions (and lower BUSCO scores) were also observed in the StringTie2 genome annotation of the water strider (*Microvelia longipes*) (Toubiana et al., 2021). In our study, *Rosa* ST2 (SR/LR) run was closest with a BUSCO score of 97.2%, BR (SR/LR) of 96.9%, and TSB (SR/ST2) of 98% (Table S9, S10).

### Including proteins: Genome annotations are improved for models,and closely related species, when full-length proteins sourced from OrthoDB are used in combination with read data

The performance of StringTie2 and Trinity-derived protein evidence was assessed on the predicted gene models using BRAKER and TSEBRA. In this context, the genome-guided or *de novo* assembled transcripts were translated into proteins and provided as evidence to train the *ab initio* component of the pipelines. Adding protein evidence to genome annotation can target protein-coding genes leading to more accurate predictions than RNA-Seq evidence alone (Bruna, 2022). This study specifically focused on using protein evidence derived in some fashion from the transcriptomic inputs but also evaluated the recommendation to include clade specific OrthoDB protein inputs to the BRAKER/TSEBRA approach.

The TSEBRA runs of the model species, *Arabidopsis* and *Populus* were compared to the reference annotations. These runs were the best for the model species in terms of sensitivity and specificity as compared to the MAKER, StringTie2, Trinity and BRAKER runs (Fig 3B). The model genomes also had very similar BUSCO completeness scores but had different mono: multi ratios with the addition of protein evidence. As expected, the model genomes benefitted the most from the inclusion of the external OrthoDB proteins in terms of annotation rate and BUSCO score (both the *Arabidopsis* and *Populus* reference proteins are contained within this resource). However, mono: multi ratio challenges remained consistent across the TSEBRA runs with varying inputs.

In the case of non-model plant genomes, TSEBRA contributed to higher mono:multi ratios, and this was very evident in *Liriodendron*. However, the BUSCO scores of the non-protein runs were lower across all runs of *Liriodendron*, compared to the protein runs. The *Rosa* TSB (SR/OrthoDB) reported the highest BUSCO score across all runs; given the phylogenetic placement of *Rosa* in comparison to *Arabidopsis*, we believe that the addition of OrthoDB greatly influenced this result. On the other hand, the annotation rate of TSB (SR/OrthoDB) remained like the other runs within *Rosa*. The higher quality of the *Rosa* genome assembly, compared to the other two non-models, could also influence the utility of the protein evidence. However, mono: multi ratios remained to be high, and annotation rates were like those of the runs without OrthoDB proteins.

TSEBRA runs with proteins sourced from genome-guided predictions perform similarly, but had lower BUSCO, higher mono: multi ratios, and total gene number when compared to the short-read only runs (Fig 4A, Table S9). Among TSEBRA runs, Trinity fares better only for *Liriodendron*, which could indicate that genome-guided proteins are not a suitable choice for a more repetitive genome. This is consistent in independent assessments between *de novo* transcriptome assemblers and genome-guided assemblers with complex genomes with fragmented genome assemblies (for example in *Ae. albopictus* (Huang et al., 2016). TSEBRA with full length proteins sourced from OrthoDB had lower BUSCO scores when compared to BR(SR) for the non-model species, *Liriodendron* and *Funaria*. In both cases, TSB (SR/OrthoDB).

The total number of genes predicted by TSEBRA and BRAKER runs remained largely the same across all species (Table S9). However, the number of mono-exonic genes increased, whereas the multi-exonic genes decreased across all TSEBRA runs in comparison to the BRAKER runs without proteins across all species. The average gene lengths also decreased, while average lengths of mono-exonics remained the same, and the lengths of multi-exonics are higher.

Initial examination of the EnTAP reciprocal BLAST assessment revealed high annotation rates for the non-model species when protein evidence was included, particularly the multi-exonics (whereas the mono-exonic percentage remained the same) (Table S9). However, this increase in multi-exonic annotation proved to be an artifact since the total number of multi-exonic models was reduced. Direct comparison of the predictions revealed that 40%-52% of the multi-exonics were split into mono-exonic predictions when comparing the BR(SR) to the TSB (SR/ST2) and TSB (SR/OrthoDB) gene models predicted using *Liriodendron* (Table S14).

### Long-read transcriptomes: Long-reads can be paired with short-reads to improve the quality of the resulting models

Long-reads generated from platforms such as Oxford Nanopore or PacBio have the potential to resolve splice variants and assemble transcripts more accurately than traditional Illumina RNA-Seq (Amarasinghe et al., 2020). While long-reads can independently generate transcriptomes, it is recommended to have a combination of short and long reads to achieve greater depth, improved error profiles, and gain more evidence for splice site resolution (Amarasinghe et al., 2020; Gonzalez-Ibeas et al., 2016; Watson & Warr, 2019).

In this study, we utilized both ONT and Iso-Seq long-reads. In the latter, we relied on raw reads (not the error-corrected CCS reads) in our comparisons for genome annotations using long-reads. In all cases, long-reads (alone) did not outperform short-reads for the BRAKER runs. However, in some cases, the combination of short-read and long-read inputs was beneficial. The higher error rate Iso-Seq reads from *Populus* and *Liriodendron* produced comparable, but lower, BUSCO scores compared to the BR (SR) runs. In contrast, the ONT long-reads used for *Arabidopsis* and *Rosa* in the combined runs (BR (SR/LR)) had slightly better BUSCO completeness as compared to the BR (SR) runs, and similar mono:multi ratios. Overall, the lower error profile of using ONT reads, supplemented with short-read data, as well as using high quality reference genomes, support the higher BUSCO completeness scores.

### Best Practices for Plant Genome Annotation

Given existing tools, we recommend that investigators utilize RepeatModeler2 to mask their genome of interest with the default settings (Flynn et al. 2020). Following soft-masking, RNA-Seq short reads (between 4-10 libraries, paired-end, minimum 15M reads per library) are generally sufficient for annotation. While we did not comprehensively investigate the impact of tissue type, it is recommended to sample from multiple tissues when possible (Kress et al., 2022). In our study, we did not observe a difference in the annotation completeness among species with a higher number of short-read libraries, although we did not comprehensively evaluate the difference of using fewer libraries within a single species.

Sequencing of long-read libraries remains more expensive than generating deep Illumina short-read RNA-Seq. In most cases, the short reads were sufficient as input. The notable exceptions include the BR (SR/LR), as they were comparable, and in some cases, slightly improved, over BR (SR) across all species. The lower error-profile Nanopore reads were more beneficial when combined with short-reads. However, current long-read technologies available from both platforms may provide different results.

BRAKER and TSEBRA outperformed runs of MAKER, StringTie2 and Trinity with default settings. It should be noted that the authors did not comprehensively benchmark MAKER with more than two training runs of AUGUSTUS as recommended, which could have further improved results. However, previous benchmarking studies also support lower performance of MAKER (Banerjee et al., 2021; Hoff et al., 2020). Among the BRAKER runs executed in the model plants, *Arabidopsis* and *Populus*, the TSEBRA runs were the best runs. TSEBRA, especially TSB (SR/OrthoDB), performs the best for *Rosa* but would require substantial filtering to remove false positives. Among the less contiguous and more evolutionary distant species (*Funaria* and *Liriodendron*), BR (SR) runs performed the best in terms of BUSCO completeness and mono:multi ratios. Overall EnTAP annotation rates were greatly improved in runs where OrthoDB proteins were included as evidence. However, when considering BUSCO as well as mono: multi ratio, especially for non-model species, the BR(SR) runs performed the best. For more divergent species (as defined by current public databases), BRAKER runs with short-reads, or short-reads and long-reads, are advised.

Regardless of approach, existing pipelines do not provide appropriate summary statistics or robust methods for filtering unlikely gene models. All methods produce more putative false positives than desired. We recommend utilizing reciprocal BLAST searches with well curated databases containing targets with full-length proteins (such as NCBI’s RefSeq) to identify fragmented models. We also recommend filtering and removing mono-exonics that do not have a protein domain. Finally, we recommend structural filters to remove unlikely gene structures (splice sites, start sites, incompletes, etc).

In this study, we demonstrated the impact of post-filtering on the most complex genome assessed in this study, *Liriodendron*. We improved the published annotation across all benchmarks evaluated in this study following a new BR(SR) run (Table 4; (Chen et al., 2019)). The filters reduced the overall number of putative false positives and increased the overall rate of annotation, with minimal reduction to BUSCO completeness.

## Data Availability

All scripts and data used is available through https://www.protocols.io/blind/3A33C8E3B76511EC84CA0A58A9FEAC02. The public data (NCBI SRA and genome assembly accessions) for the reference genomes, short-reads, and long-reads are listed in Table S2.

## Acknowledgements

The authors would like to thank the Institute for Systems Genomics (ISG) and Computational Biology Core at UConn for HPC services. Thanks go to Dr. Bernard Goffinet for his careful review of this manuscript. This work was supported by the National Science Foundation awards, DEB-1753811 and DBI-1943371.

