## Supplement Table 1 for "Welcome to the big leaves: best practices for improving genome annotation in non-model plant genomes"

*Table S1: Genomes for this study, with versions and links to the genome*

| **Species** | **Paper** | **Version** | **Link to Genome** |
|---|---|---|---|
| *Arabidopsis thaliana* | (Lamesch et al. 2012) | TAIR10 - release 41 | ftp://ftp.ensemblgenomes.org/pub/plants/release-41/fasta/arabidopsis_thaliana/dna/Arabidopsis_thaliana.TAIR10.dna_sm.toplevel.fa.gz |
| *Funaria hygrometrica* | (Kirbis et al. 2022) | Version 1 | Inhouse |
| *Populus trichocarpa* | (Tuskan et al. 2006) | Version 3 | ftp://ftp.ensemblgenomes.org/pub/plants/release-49/fasta/populus_trichocarpa/dna/ |
| *Liriodendron chinense* | (Chen et al. 2019) | Version 1 | https://hardwoodgenomics.org/sites/default/files/sequences/liriodendron_chinense/LIR.pbjelly.reN.final.fasta |
| *Rosa chinensis* | (Raymond et al. 2018) | Version 1 | https://www.rosaceae.org/rosaceae_downloads/Rosa_chinensis/Rchinensis_Old_Blush_homozygous_genome-v2.0/assembly/Rosa_chinensis_Old_Blush_homozygous_genome-v2.0.fna.gz |


Chen, Jinhui, Zhaodong Hao, Xuanmin Guang, Chenxi Zhao, Pengkai Wang, Liangjiao Xue, Qihui Zhu, et al. 2019. “Liriodendron Genome Sheds Light on Angiosperm Phylogeny and Species-Pair Differentiation.” *Nature Plants* 5 (1): 18–25.

Kirbis, Alexander, Nasim Rahmatpour, Shanshan Dong, Jin Yu, Nico van Gessel, Manuel Waller, Ralf Reski, et al. 2022. “Genome Dynamics in Mosses: Extensive Synteny Coexists with a Highly Dynamic Gene Space.” *bioRxiv*. https://doi.org/10.1101/2022.05.17.492078.

Lamesch, Philippe, Tanya Z. Berardini, Donghui Li, David Swarbreck, Christopher Wilks, Rajkumar Sasidharan, Robert Muller, et al. 2012. “The Arabidopsis Information Resource (TAIR): Improved Gene Annotation and New Tools.” *Nucleic Acids Research* 40 (Database issue): D1202–10.

Raymond, Olivier, Jérôme Gouzy, Jérémy Just, Hélène Badouin, Marion Verdenaud, Arnaud Lemainque, Philippe Vergne, et al. 2018. “The Rosa Genome Provides New Insights into the Domestication of Modern Roses.” *Nature Genetics* 50 (6): 772–77.

Tuskan, G. A., S. Difazio, S. Jansson, J. Bohlmann, I. Grigoriev, U. Hellsten, N. Putnam, et al. 2006. “The Genome of Black Cottonwood, Populus Trichocarpa (Torr. & Gray).” *Science* 313 (5793): 1596–1604.

*Table S2: Public transcriptomic evidence (short and long read)*

| Species | | Bioproject # | SRA (SRP/ERP) Study | SRX/ERX Experiment | SRR/ERR Run | Spots (# read locations) | Bases | Bytes | Tissue type | Sequencer | Paired/single |
|---|---|---|---|---|---|---|---|---|---|---|---|
| ***Funaria hygrometrica*** | | PRJNA421369 | SRP126294 | SRX3452863 | SRR8527271 | 49,324,732 | 7.4G | 2.5Gb | Haploid gametophyte | Illumina NextSeq 500 | paired |
| | | | | SRX5330016 | SRR6356272 | 34,002,818 | 5.1G | 1.7Gb | Haploid gametophyte | Illumina NextSeq 500 | paired |
| | | | | SRX5330017 | SRR8527272 | 37,100,201 | 5.6G | 1.9Gb | Haploid gametophyte | Illumina NextSeq 500 | paired |
| ***Arabidopsis thaliana*** | | PRJNA284739 | SRP058628 | SRX1036705 | SRR2037363 | 82,831,218 | 16.6G | 10.6Gb | Stem | Illumina HiSeq 2000 | paired |
| | | PRJNA284739 | SRP058628 | SRX1036705 | SRR2037362 | 39,299,841 | 7.9G | 5.1Gb | Stem | Illumina HiSeq 2000 | paired |
| | | PRJNA284739 | SRP058628 | SRX1036693 | SRR2037338 | 57,633,045 | 11.5G | 7.5Gb | Stem | Illumina HiSeq 2000 | paired |
| | | PRJNA284739 | SRP058628 | SRX1036695 | SRR2037346 | 64,896,842 | 13G | 8.5Gb | Leaf | Illumina HiSeq 2000 | paired |
| | | PRJEB14117 | ERP015730 | ERX2770903 | ERR2757910 | 36,319,987 | 10.3G | 4Gb | Leaf | Illumina HiSeq 2000 | paired |
| | | PRJNA340889 | SRP083317 | SRX2065453 | SRR4095612 | 17,676,726 | 3.6G | 1.5Gb | Seeds | Illumina HiSeq 2500 | paired |
| | | PRJNA340889 | SRP083317 | SRX2065448 | SRR4095607 | 14,959,896 | 3G | 1.3Gb | Seeds | Illumina HiSeq 2500 | paired |
| | | PRJEB42556 | ERP126434 | ERX5132034 | ERR5343646 | 10,151,920 | 3G | 1.2Gb | Aerial Part | Illumina HiSeq 3000 | paired |

| | | | | | | | | | | | |
|---|---|---|---|---|---|---|---|---|---|---|---|
| | | PRJEB40453 | ERP124098 | ERX4548377 | ERR4619338 | 24,106,303 | 7.3G | 3Gb | 17 day-old shoots | Illumina HiSeq 3000 | paired |
| | | PRJNA594286 | SRP235227 | SRX7290465 | SRR10611193 | 8,258,511 | 10.1G | 8Gb | Unkown | Oxford Nanopore PromethION | single |
| | | PRJNA594286 | SRP235227 | SRX7290466 | SRR10611194 | 7,849,574 | 9.8G | 7.8Gb | Unkown | Oxford Nanopore PromethION | single |
| | | PRJNA594286 | SRP235227 | SRX7290467 | SRR10611195 | 7,025,983 | 8.6G | 6.8Gb | Unkown | Oxford Nanopore PromethION | single |
| | | PRJNA725066 | SRP316288 | SRX10677446 | SRR14322310 | 68,785,614 | 20.6G | 6.1Gb | axillary buds | Illumina NovaSeq 6000 | paired |
| ***Populus trichocarpa*** | | PRJNA515420 | SRP179723 | SRX9893893 | SRR13481183 | 21,263,642 | 5.3G | 1.8Gb | Secondary xylem | Illumina HiSeq 2500 | paired |
| | | PRJNA628501 | SRP258577 | SRX8177635 | SRR11611372 | 13,991,070 | 4.2G | 1.4Gb | seedling needles | Illumina HiSeq 2500 | paired |
| | | PRJNA650996 | SRP289313 | SRX9383942 | SRR12919313 | 22,476,947 | 6.8G | 2Gb | bark | Illumina NovaSeq 6000 | paired |
| | | PRJNA696790 | SRP306056 | SRX10084749 | SRR13695406 | 20,039,851 | 6.1G | 1.8Gb | leaf | Illumina NovaSeq 6000 | paired |
| | | PRJNA725066 | SRP316288 | SRX10677446 | SRR14322310 | 68,785,614 | 20.6G | 6.1Gb | axillary buds | Illumina NovaSeq 6000 | paired |
| | | PRJNA516416 | SRP179723 | SRX5254305 | SRR8447264 | 161,334 | 383.7M | 93.1Mb | secondary xylem | PacBio SMRT | single |
| ***Liriodendron chinense*** | | PRJNA559687 | SRP218024 | SRX6693922 | SRR9945429 | 45,919,256 | 13.8G | 4Gb | bract | Illumina HiSeq 2500 | paired |

| | | | | | | | | | | | |
|---|---|---|---|---|---|---|---|---|---|---|---|
| | | | | SRX6697492 | SRR9949010 | 38,265,327 | 11.5G | 3.3Gb | bract | Illumina HiSeq 2500 | paired |
| | | | | SRX6693921 | SRR9945430 | 39,896,641 | 12G | 3.4Gb | leaf | Illumina HiSeq 2500 | paired |
| | | | | SRX6697491 | SRR9949011 | 34,672,556 | 10.4G | 2.9Gb | leaf | Illumina HiSeq 2500 | paired |
| | | | | SRX6697402 | SRR9948916 | 41,436,299 | 12.4G | 3.6Gb | leaf | Illumina HiSeq 2500 | paired |
| | | | | SRX6693920 | SRR9945431 | 38,819,287 | 11.6G | 3.4Gb | petal | Illumina HiSeq 2500 | paired |
| | | | | SRX6697494 | SRR9949008 | 29,694,072 | 8G | 2.3Gb | petal | Illumina HiSeq 2500 | paired |
| | | | | SRX6697399 | SRR9948919 | 37,405,592 | 11.2G | 3.2Gb | petal | Illumina HiSeq 2500 | paired |
| | | | | SRX6693919 | SRR9945432 | 30,029,905 | 9G | 2.6Gb | pistil | Illumina HiSeq 2500 | paired |
| | | | | SRX6697493 | SRR9949009 | 41,125,315 | 12.3G | 3.5Gb | pistil | Illumina HiSeq 2500 | paired |
| | | | | SRX6697400 | SRR9948918 | 34,728,258 | 10.4G | 3Gb | pistil | Illumina HiSeq 2500 | paired |
| | | | | SRX6693918 | SRR9945433 | 44,447,597 | 13.3G | 3.8Gb | shoot apex | Illumina HiSeq 2500 | paired |
| | | | | SRX6697496 | SRR9949006 | 37,881,676 | 11.4G | 3.3Gb | shoot apex | Illumina HiSeq 2500 | paired |
| | | | | SRX6697404 | SRR9948914 | 37,425,347 | 11.2G | 3.2Gb | shoot apex | Illumina HiSeq 2500 | paired |

| | | | | | | | | | | | |
|---|---|---|---|---|---|---|---|---|---|---|---|
| | | | | SRX6693917 | SRR9945434 | 45,653,579 | 13.7G | 3.9Gb | sepal | Illumina HiSeq 2500 | paired |
| | | | | SRX6697495 | SRR9949007 | 43,209,095 | 13G | 3.7Gb | sepal | Illumina HiSeq 2500 | paired |
| | | | | SRX6697405 | SRR9948913 | 33,151,069 | 9.9G | 2.8Gb | sepal | Illumina HiSeq 2500 | paired |
| | | | | SRX6693916 | SRR9945435 | 46,620,902 | 14G | 4Gb | stamen | Illumina HiSeq 2500 | paired |
| | | | | SRX6697497 | SRR9949005 | 38,971,271 | 11.7G | 3.3Gb | stamen | Illumina HiSeq 2500 | paired |
| | | | | SRX6697403 | SRR9948915 | 41,065,829 | 12.3G | 3.5Gb | stamen | Illumina HiSeq 2500 | paired |
| | | PRJNA665077 | SRP285036 | SRX9174881 | SRR12695187 | 10,437,029 | 22.7G | 5.4Gb | All 7 samples above pooled together | PACBIO-SMRT | single |
| ***Rosa chinensis*** | | PRJNA398090 | SRP115334 | SRX6490239 | SRR9733228 | 37,170,059 | 11.2G | 3.2Gb | the abaxial side of petal | Illumina HiSeq 4000 | paired |
| | | PRJNA473465 | SRP150297 | SRX4195946 | SRR7293274 | 35,120,535 | 10.5G | 3.9Gb | leaves | Illumina HiSeq 2000 | paired |
| | | PRJNA473465 | SRP150297 | SRX4195940 | SRR7293280 | 33,483,963 | 10G | 3.8Gb | leaves | Illumina HiSeq 2000 | paired |
| | | PRJNA546486 | SRP200448 | SRX5973706 | SRR9202384 | 30,805,477 | 9.2G | 2.7Gb | Stamen | Illumina HiSeq 4000 | paired |
| | | PRJNA546486 | SRP200448 | SRX5973701 | SRR9202389 | 29,793,315 | 8.9G | 2.7Gb | Pistil_Ovary | Illumina HiSeq 4000 | paired |

| | | | | | | | | | | | |
|---|---|---|---|---|---|---|---|---|---|---|---|
| | | PRJNA546486 | SRP200448 | SRX5973705 | SRR9202385 | 28,535,324 | 8.6G | 2.5Gb | Prickle | Illumina HiSeq 4000 | paired |
| | | PRJNA596583 | SRP238318 | SRX7419418 | SRR10744004 | 22,985,180 | 6.8G | 2Gb | leaf | Illumina HiSeq 4000 | paired |
| | | PRJNA414720 | SRP120300 | SRX3296708 | SRR6186662 | 23,141,140 | 6.7G | 2.3Gb | leaf | Illumina HiSeq 2500 | paired |
| | | PRJNA414720 | SRP120300 | SRX3296710 | SRR6186660 | 27,768,588 | 5G | 1.9Gb | shoot apical meristem | Illumina HiSeq 2500 | paired |
| | | PRJNA236618 | SRP035933 | SRX451091 | SRR1145848 | 33,261,614 | 4.8G | 2.6Gb | Flower bud, open flower and senescent flower | Illumina HiSeq 2000 | paired |
| | | | | SRX9153921 | SRR12673743 | 7,290,141 | 8.2G | 6.9Gb | | | |
| | | | | SRX9153918 | SRR12673746 | 6,924,721 | 8.7G | 7.3Gb | | | |
| | | | | SRX9153916 | SRR12673748 | 6,747,461 | 7G | 5.9Gb | | | |
| | | | | SRX9153914 | SRR12673750 | 6,585,762 | 7.3G | 6.1Gb | | | |
| | | | | SRX9153912 | SRR12673752 | 7,641,480 | 9.1G | 7.6Gb | | | |
| | | | | SRX9153909 | SRR12673755 | 6,739,818 | 8G | 6.7Gb | | | |

*Table S3: Mono: multi ratios in published model plant genomes*

| Species | Mono | Multi | Total | Ratio | Source |
|---|---|---|---|---|---|
| **Arabidopsis** | 8093 | 40227 | 48320 | 0.20 | ftp://ftp.ensemblgenomes.org/pub/plants/release-41/fasta/arabidopsis_thaliana/dna/Arabidopsis_thaliana.TAIR10.dna_sm.toplevel.fa.gz |
| **Populus** | 10802 | 62210 | 73012 | 0.17 | ftp://ftp.ensemblgenomes.org/pub/plants/release-49/fasta/populus_trichocarpa/dna/ |
| ***Oryza sativa*** | 8825 | 33715 | 42540 | 0.26 | https://ftp.ncbi.nlm.nih.gov/genomes/all/GCF/001/433/935/GCF_001433935.1_IRGSP-1.0/GCF_001433935.1_IRGSP-1.0_genomic.fna.gz |
| ***Solanum lycopersicum*** | 6123 | 31460 | 37583 | 0.19 | https://ftp.ncbi.nlm.nih.gov/genomes/all/GCF/000/188/115/GCF_000188115.5_SL3.1/GCF_000188115.5_SL3.1_genomic.fna.gz |
| ***Nicotiana tabacum*** | 15220 | 69003 | 84223 | 0.22 | https://ftp.ncbi.nlm.nih.gov/genomes/all/GCF/000/715/135/GCF_000715135.1_Ntab-TN90/GCF_000715135.1_Ntab-TN90_genomic.fna.gz |

*Table S4: Initial genome statistics*

| Species | | *Arabidopsis* | *Funaria* | *Liriodendron* | *Populus* | *Rosa* |
|---|---|---|---|---|---|---|
| **Genome size (bp)** | | 119,667,750 | 326,856,579 | 1,742,423,874 | 434,132,815 | 515,588,973 |
| **# of contigs** | | 7 | 687 | 3711 | 1446 | 55 |
| **N50** | | 23,459,830 | 1,484,274 | 3,525,943 | 19,465,461 | 69,643,165 |
| **Repeat content** | | 23.6 | 42.35 | 73.18 | 35.9 | cd |
| **Repeat content (RM2+)** | | 16.51 | 43.12 | 72.66 | 45.06 | |
| **BUSCO (genome)** | | C:99.3%[S:98.6%, D:0.7%],F:0.2%,M:0.5%,n:1614 | C:85.6%[S:73.5%,D:12.1%],F:2.4%,M:12.0%,n:1614 | C:98.6%[S:92.1%,D:6.5%],F:0.8%,M:0.6%,n:161 | C:98.8%[S:80.6%,D:18.2%], F:0.6%,M:0.6%,n:1614 | C:98.8%[S:94.4%,D:4.4%],F:0.6%,M:0.6%,n:1614 |
| **BUSCO (annotated proteins)** | | C:99.6%[S:55.8%, D:43.8%],F:0.1%, M:0.3%,n:161 | C:86.6%[S:72.6%,D:14.0%],F:2.3%,M:11.1%,n:1614 | C:75.1%[S:68.3%,D:6.8%],F:15.1%,M:9.8%,n:1614 | C:98.3%[S:35.0%,D:63.3%], F:0.9%,M:0.8%,n:1614 | C:97.3%[S:93.4%,D:3.9%],F:1.7%,M:1.0%,n:1614 |
| **gFACs (reference annotation)** | **Mono** | 8093 | 15640 | 14521 | 10802 | 15383 |
| | **Multi** | 40227 | 20620 | 20740 | 62210 | 35004 |
| | **Total** | 48320 | 36260 | 35261 | 73012 | 50387 |
| | **Ratio** | 0.20 | 0.76 | 0.70 | 0.17 | 0.44 |

*Table S5: RNA alignments for short and long reads*

| **Reference genome** | **RNA read alignments, reference species (%)** | | ***De novo* Transcripts aligned to genome** | |
|---|---|---|---|---|
| | **Short-reads** | **Long-reads (minimap2-sequencer)** | **(% aligned) mono/multi** | **BUSCO Completeness** |
| *Funaria* | 97.13 | N/A | [94.3] 15179/39149 | C:57.2%[S:43.6%,D:13.6%],F:13.6%,M:29.2%,n:1614 |
| *Arabidopsis* | 97.06 | 97.1 (ONT) | [94.7] 11291/34492 | C:78.8%[S:67.5%,D:11.3%],F:13.6%,M:7.6%,n:1614 |
| *Populus* | 91.41 | 92.01 (PacBio) | [73.7] 21604/57170 | C:78.7%[S:52.0%,D:26.7%],F:12.7%,M:8.6%,n:1614 |
| *Liriodendron* | 97.05 | 95.5 (PacBio) | [69.6] 66629/79698 | C:76.5%[S:47.1%,D:29.4%],F:13.8%,M:9.7%,n:1614 |
| *Rosa* | 91.96 | 99 (ONT) | [72.8] 29985/65031 | C:83.2%[S:57.4%,D:25.8%],F:12.0%,M:4.8 |

| | | | | |
|---|---|---|---|---|
| | | | | %,n:1614 |

*Table S6: N50s for the SR and LR reads*

| Species | SR- Total reads (N50) | LR- Total reads (N50) |
|---|---|---|
| ***Arabidopsis*** | 52,912,443,562 (100) | 72,435,151,489 (2249) |
| ***Liriodendron*** | 208,110,232,535 (150) | 22,713,962,011 (4673) |
| ***Populus*** | 53,390,254,188 (150) | 558,593,446 (1,348) |
| ***Rosa*** | 135,933,621,273 (150) | 482,704,44,525 (976) |
| ***Funaria*** | 43,506,430,892 (76) | |

*Table S7: Sensitivity and Precision scores for Arabidopsis and Populus*

| Species | Run | Sensitivity | Precision |
|---|---|---|---|
| **Arabidopsis** | MK (RM2+) | 6.41 | 1.85 |
| | BR (SR) | 40.37 | 60.11 |
| | TSB (SR/TRINITY) | 55.94 | 80.78 |
| | TSB (SR/ST2) | 56.94 | 81.92 |
| | BR (SR/LR) | 42.45 | 61.88 |
| | TSB (SR/LR/ST2) | 56.21 | 82.40 |
| | BR (LR) | 42.46 | 59.79 |
| | TSB (LR/ST2) | 54.91 | 81.17 |
| | BR (SR/RM2+) | 43.67 | 61.70 |
| | TSB (SR/RM2+/ST2) | 56.99 | 81.93 |
| | ST2 (SR) | 50.07 | 63.24 |
| | ST2 (LR/SR) | 46.41 | 51.99 |
| | ST2 (LR) | 39.57 | 50.14 |
| | TSB (SR/OrthoDB) | 58.5 | 80.85 |
| **Populus** | MK (RM2+) | 0.40 | 0.31 |

| | | | |
|---|---|---|---|
| | BR (SR) | 32.01 | 41.02 |
| | TSB (SR/TRINITY) | 39.50 | 62.58 |
| | TSB (SR/ST2) | 41.83 | 62.77 |
| | BR (SR/LR) | 31.87 | 40.79 |
| | TSB (SR/LR/ST2) | 41.55 | 62.67 |
| | BR (LR) | 32.45 | 42.23 |
| | TSB (LR/ST2) | 39.85 | 62.82 |
| | BR (SR/RM2+) | 31.73 | 41.25 |
| | TSB (SR/RM2+/ST) | 41.55 | 60.56 |
| | ST2 (SR) | 29.86 | 45.43 |
| | ST2 (LR/SR) | 13.08 | 56.89 |
| | ST2 (LR) | 19.96 | 56.68 |
| | TSB (SR/OrthoDB) | 40.28 | 57.24 |

*Table S8: Gene characteristics in BRAKER runs*

| | | | Gene length | | |
|---|---|---|---|---|---|
| **Species** | **Runs** | **Average gene length** | **mono** | **multi** | **Median number of exons per multiexonic gene:** |
| *Arabidopsis* | **BR (SR)** | 2936.07 | 854.67 | 3408.93 | 5 |
| | **BR (LR)** | 2229.38 | 836.10 | 2569.73 | 5 |
| | **BR (SR/LR)** | 2059.37 | 849.09 | 2338.11 | 5 |
| | **TSB (SR/ST2)** | 1860.28 | 876.37 | 2245.72 | 5 |
| | **TSB (SR/TRINITY)** | 1860.28 | 876.37 | 2245.72 | 5 |
| | **TSB (LR/ST2)** | 1945.61 | 881.37 | 2348.70 | 5 |
| | **TSB (SR/LR/ST2)** | 1957.18 | 885.25 | 2364.78 | 5 |
| | **BR (SR/RM2+)** | 3629.76 | 892.27 | 4233.12 | 5 |
| | **TSB** | 1942.28 | 887.28 | 2355.87 | 5 |

| | | | | | |
|---|---|---|---|---|---|
| | **(SR/ST2/RM2+)** | | | | |
| | **TSB (SR/OrthoDB)** | 1929.71 | 877.52 | 2289.58 | 5 |
| *Funaria* | **BR (SR)** | 2112.89 | 592.61 | 2734.37 | 4 |
| | **TSB (SR/ST2)** | 1701.69 | 627.82 | 3008.18 | 5 |
| | **TSB (SR/TRINITY)** | 2061.61 | 586.63 | 3304.85 | 6 |
| | **BR (SR/RM2+)** | 2124.45 | 604.88 | 2722.5 | 4 |
| | **TSB (SR/ST2/RM2+)** | 1702.21 | 624.92 | 3007.26 | 5 |
| | **TSB (SR/OrthoDB)** | 2072.513 | 584.954 | 3300.495 | 6 |
| *Liriodendron* | **BR (SR)** | 9399.97 | 682.3 | 12415.27 | 4 |
| | **BR (LR)** | 8601.03 | 675.74 | 12149.00 | 4 |
| | **BR (SR/LR)** | 9725.47 | 694.18 | 12734.64 | 4 |
| | **TSB (SR/ST2)** | 6596.95 | 734.05 | 12274.35 | 5 |
| | **TSB (SR/TRINITY)** | 6229.90 | 724.99 | 11872.86 | 4 |
| | **TSB (LR/ST2)** | 6409.38 | 737.48 | 12204.38 | 4 |
| | **BR (SR/RM2+)** | 9268.75 | 654.36 | 12326.227 | 4 |
| | **TSB (SR/LR/ST2)** | 6238.68 | 733.26 | 11909.92 | 4 |
| | **TSB (SR/RM2+/ST2)** | 6271.92 | 718.36 | 12040.91 | 4 |
| | **TSB (SR/OrthoDB)** | 6886.243 | 732.22 | 12954.44 | 5 |
| *Populus* | **BR (SR)** | 3142.98 | 800.94 | 3709.66 | 4 |
| | **BR (LR)** | 6963.77 | 792.77 | 8734.16 | 4 |
| | **BR (SR/LR)** | 3970.86 | 748.12 | 4859.46 | 4 |
| | **TSB (SR/ST2)** | 2724.53 | 841.97 | 3610.33 | 4 |
| | **TSB (SR/TRINITY)** | 2764.77 | 831.73 | 3706.87 | 4 |
| | **TSB (LR/ST2)** | 2703.65 | 818.37 | 3641.62 | 4 |
| | **TSB (SR/LR/ST2)** | 2719.63 | 843.70 | 3599.58 | 4 |
| | **BR (SR/RM2+)** | 4596.54 | 735.48 | 5678.38 | 4 |
| | **BR (SR/ST2/RM2+)** | 2648.45 | 784.78 | 3593.29 | 4 |
| | **TSB (SR/OrthoDB)** | 2864.61 | 796.04 | 3590.612 | 5 |
| *Rosa* | **BR (SR)** | 2372.42 | 723.95 | 2993.17 | 4 |
| | **BR (LR)** | 3765.28 | 752.45 | 4952.92 | 4 |
| | **BR (SR/LR)** | 3977.25 | 712.35 | 5289.26 | 4 |
| | **TSB (SR/ST2)** | 2064.18 | 777.14 | 3033.22 | 4 |
| | **TSB (SR/TRINITY)** | 2037.87 | 766.38 | 3010.40 | 4 |
| | **TSB (LR/ST2)** | 3765.28 | 752.45 | 4952.92 | 4 |
| | **TSB (SR/LR/ST2)** | 2059.58 | 791.41 | 3012.58 | 4 |

| | | | | | |
|---|---|---|---|---|---|
| | **TSB (SR/OrthoDB)** | 2065.471 | 77.69 | 2997.96 | 5 |

*Table S9: Comparisons among Stringtie2,  BRAKER, and TSEBRA runs*

| ***Species*** | **Runs** | **Total** | **Annotated genes (70/70)** | **Annotation rate** | **Percentage gene family assignment** | **mono** | **multi** | **ratio** | **monos annotated** | **multi annotated** | **%monos annotated** | **%multis annotated** | **busco** |
|---|---|---|---|---|---|---|---|---|---|---|---|---|---|
| ***Arabidopsis*** | BR (SR) | 27365 | 24594 | 89.87 | 0.96 | 5065 | 22296 | 0.23 | 4461 | 20129 | 88.07 | 90.28 | C:96.3%[S:90.2%,D:6.1%],F:1.0%,M:2.7%,n:1614 |
| | BR (LR) | 28469 | 25512 | 89.61 | 0.96 | 5589 | 22880 | 0.24 | 4812 | 20700 | 86.10 | 90.47 | C:97.1%[S:87.3%,D:9.8%],F:1.1%,M:1.8%,n:1614 |
| | BR (SR/LR) | 27826 | 24960 | 89.70 | 0.96 | 5209 | 22617 | 0.23 | 4589 | 20363 | 88.10 | 90.03 | C:97.3%[S:90.9%,D:6.4%],F:0.8%,M:1.9%,n:1614 |
| | TSB (SR/ST2) | 27178 | 25619 | 94.26 | 0.98 | 7515 | 19663 | 0.38 | 6569 | 19050 | 87.42 | 96.89 | C:98.6%[S:90.7%,D:7.9%],F:0.2%,M:1.2%,n:1614 |
| | TSB (SR/TRINITY) | 27162 | 25546 | 94.05 | 0.98 | 7643 | 19519 | 0.39 | 6618 | 18923 | 86.56 | 96.95 | C:97.7%[S:90.9%,D:6.8%],F:0.9%,M:1.4%,n:1614 |
| | TSB (LR/ST2) | 26373 | 24784 | 93.97 | 0.96 | 7245 | 19128 | 0.38 | 6245 | 18539 | 86.20 | 96.92 | C:96.8%[S:90.1%,D:6.7%],F:1.1%,M:2.1%,n:1614 |
| | TSB (SR/LR/ST2) | 26545 | 24990 | 94.14 | 0.98 | 7313 | 19232 | 0.38 | 6329 | 18661 | 86.54 | 97.03 | C:98.1%[S:90.9%,D:7.2%],F:0.4%,M:1.5%,n:1614 |
| | BR (RM2/SR) | 28660 | 25205 | 87.94 | 0.94 | 5176 | 23484 | 0.22 | 4557 | 20645 | 88.04 | 87.91 | C:97.3%[S:90.6%,D:6.7%],F:0.9%,M:1.8%,n:1614 |
| | TSB (SR/RM2/ST2) | 27185 | 25505 | 93.82 | 0.97 | 7656 | 19529 | 0.39 | 6570 | 18935 | 85.81 | 96.96 | C:98.6%[S:91.5%,D:7.1%],F:0.3%,M:1.1%,n:1614 |
| ***Funaria*** | BR (SR) | 52000 | 22500 | 43.27 | 0.45 | 15089 | 36911 | 0.41 | 2737 | 19763 | 18.14 | 53.54 | C:85.8%[S:63.8%,D:22.0%],F:2.9%,M:11.3%,n:1614 |

| ***Species*** | **Runs** | **Total** | **Annotated genes (70/70)** | **Annotation rate** | **Percentage gene family assignment** | **mono** | **multi** | **ratio** | **monos annotated** | **multi annotated** | **%monos annotated** | **%multis annotated** | **busco** |
|---|---|---|---|---|---|---|---|---|---|---|---|---|---|
| | TSB (SR/ ST2) | 45884 | 22608 | 49.27 | 0.64 | 25184 | 20700 | 1.22 | 4867 | 17201 | 19.32 | 83.10 | C:82.2%[S:69.3%,D:12.9%],F:2.8%,M:15.0%,n:1614 |
| | TSB (SR/ TRINITY) | 31928 | 16989 | 53.21 | 0.61 | 14603 | 17325 | 0.84 | 2724 | 14265 | 18.65 | 82.34 | C:86.6%[S:66.2%,D:20.4%],F:2.2%,M:11.2%,n:1614 |
| | BR (SR/ RM2+) | 50408 | 22142 | 43.92 | 0.59 | 14236 | 36175 | 0.39 | 2651 | 19491 | 18.62 | 53.88 | C:85.9%[S:64.7%,D:21.2%],F:2.4%,M:11.7%,n:1614 |
| | TSB (SR/ RM2+/ST2) | 45792 | 22013 | 48.07 | 0.61 | 26358 | 20688 | 1.27 | 4826 | 17187 | 18.31 | 83.08 | C:84.5%[S:41.2%,D:43.3%],F:3.0%,M:12.5%,n:1614 |
| ***Liriodendron*** | BR (SR) | 52157 | 31123 | 59.67 | 0.75 | 13404 | 38755 | 0.34 | 6707 | 24417 | 50.04 | 63.00 | C:90.8%[S:72.7%,D:18.1%],F:5.9%,M:3.3%,n:1614 |
| | BR (LR) | 50343 | 30907 | 61.39 | 0.75 | 15568 | 34775 | 0.45 | 7618 | 23289 | 48.93 | 66.97 | C:88.2%[S:70.9%,D:17.3%],F:7.1%,M:4.7%,n:1614 |
| | BR (SR/ LR) | 51008 | 30709 | 60.20 | 0.75 | 12747 | 19996 | 0.64 | 6463 | 12556 | 50.70 | 62.79 | C:90.9%[S:72.1%,D:18.8%],F:5.3%,M:3.8%,n:1614 |
| | TSB (SR/ ST2) | 49150 | 33017 | 67.17 | 0.78 | 24180 | 24970 | 1.00 | 12145 | 20872 | 50.23 | 83.59 | C:84.2%[S:73.5%,D:10.7%],F:8.2%,M:7.6%,n:1614 |
| | TSB (SR/ TRINITY) | 52407 | 34427 | 65.70 | 0.77 | 26528 | 25879 | 1.02 | 13196 | 21231 | 49.74 | 82.04 | C:84.7%[S:74.2%,D:10.5%],F:7.6%,M:7.7%,n:1614 |
| | TSB (LR/ ST2) | 49740 | 33166 | 66.68 | 0.78 | 25137 | 24603 | 1.02 | 12727 | 20439 | 50.63 | 83.08 | C:82.9%[S:72.5%,D:10.4%],F:9.0%,M:8.1%,n:1614 |
| | BR (SR/ | 51788 | 31277 | 60.39 | 0.76 | 13566 | 38222 | 0.35 | 6806 | 24469 | 50.17 | 64.02 | C:88.9%[S:71.4%,D:17.5%],F:6. |

| ***Species*** | **Runs** | **Total** | **Annotated genes (70/70)** | **Annotation rate** | **Percentage gene family assignment** | **mono** | **multi** | **ratio** | **monos annotated** | **multi annotated** | **%monos annotated** | **%multis annotated** | **busco** |
|---|---|---|---|---|---|---|---|---|---|---|---|---|---|
| | RM2+) | | | | | | | | | | | | 3%,M:4.8%,n:1614 |
| | TSB (SR/LR/ST2) | 51630 | 34301 | 66.44 | 0.78 | 26198 | 25432 | 1.03 | 13205 | 21096 | 50.40 | 82.95 | C:83.2%[S:72.4%,D:10.8%],F:9.4%,M:7.4%,n:161 |
| | TSB (SR/RM2+/ST2) | 50666 | 34096 | 67.29 | 0.78 | 25815 | 24851 | 1.04 | 13320 | 20776 | 51.60 | 83.60 | C:83.6%[S:72.9%,D:10.7%],F:9.1%,M:7.3%,n:1614 |
| ***Populus*** | BR (SR) | 48424 | 36568 | 75.52 | 0.87 | 9434 | 38991 | 0.24 | 6666 | 29902 | 70.66 | 76.69 | C:97.9%[S:75.0%,D:22.9%],F:1.0%,M:1.1%,n:1614 |
| | BR (LR) | 47104 | 35851 | 76.11 | 0.87 | 10513 | 36606 | 0.29 | 7281 | 28569 | 69.26 | 78.04 | C:96.4%[S:76.0%,D:20.4%],F:2.1%,M:1.5%,n:1614 |
| | BR (SR/LR) | 48715 | 36523 | 74.97 | 0.86 | 10529 | 38188 | 0.27 | 6790 | 29733 | 64.49 | 77.86 | C:97.8%[S:75.0%,D:22.8%],F:0.9%,M:1.3%,n:1614 |
| | TSB (SR/ST2) | 40841 | 35948 | 88.02 | 0.94 | 13068 | 27773 | 0.47 | 9354 | 26594 | 71.58 | 95.75 | C:97.5%[S:72.2%,D:25.3%],F:0.7%,M:1.8%,n:1614 |
| | TSB (SR/TRINITY) | 38517 | 33686 | 87.46 | 0.93 | 12621 | 25896 | 0.49 | 8814 | 24872 | 69.83 | 96.04 | C:96.5%[S:73.3%,D:23.2%],F:1.1%,M:2.4%,n:1614 |
| | TSB (LR/ST2) | 38913 | 34061 | 87.53 | 0.93 | 12928 | 25985 | 0.50 | 9095 | 24965 | 70.35 | 96.07 | C:94.9%[S:73.2%,D:21.7%],F:2.2%,M:2.9%,n:1614 |
| | TSB (SR/LR/ST2) | 40758 | 35931 | 88.16 | 0.94 | 13014 | 27744 | 0.47 | 9336 | 26595 | 71.74 | 95.86 | C:96.1%[S:72.1%,D:24.0%],F:1.7%,M:2.2%,n:1614 |
| | BR (SR/RM2+) | 47956 | 36528 | 76.17 | 0.87 | 10494 | 37461 | 0.28 | 6717 | 29803 | 64.01 | 79.56 | C:98.0%[S:74.8%,D:23.2%],F:0.8%,M:1.2%,n:1614 |

| ***Species*** | **Runs** | **Total** | **Annotated genes (70/70)** | **Annotation rate** | **Percentage gene family assignment** | **mono** | **multi** | **ratio** | **monos annotated** | **multi annotated** | **%monos annotated** | **%multis annotated** | **busco** |
|---|---|---|---|---|---|---|---|---|---|---|---|---|---|
| | BR (SR/RM2+/ST2) | 42093 | 36016 | 85.56 | 0.92 | 14161 | 27932 | 0.51 | 9366 | 26650 | 66.14 | 95.41 | C:96.5%[S:73.4%,D:23.1%],F:1.7%,M:1.8%,n:1614 |
| ***Rosa*** | BR (SR) | 47318 | 32248 | 68.15 | 0.81 | 12944 | 34370 | 0.38 | 7275 | 24970 | 56.20 | 72.65 | C:96.4%[S:86.8%,D:9.6%],F:1.9%,M:1.7%,n:1614 |
| | BR (LR) | 47411 | 26290 | 55.45 | 0.81 | 13407 | 34006 | 0.39 | 6429 | 19861 | 47.95 | 58.40 | C:82.9%[S:79.4%,D:3.5%],F:1.9%,M:15.2%,n:1614 |
| | BR (SR/LR) | 49128 | 33002 | 67.17 | 0.80 | 14077 | 35046 | 0.40 | 7642 | 25356 | 54.29 | 72.35 | C:96.9%[S:86.9%,D:10.0%],F:1.6%,M:1.5%,n:1614 |
| | TSB (SR/ST2) | 44577 | 33814 | 75.85 | 0.85 | 19147 | 25430 | 0.75 | 11144 | 22670 | 58.20 | 89.15 | C:98.0%[S:86.4%,D:11.6%],F:0.4%,M:1.6%,n:1614 |
| | TSB (SR/TRINITY) | 45971 | 34424 | 74.88 | 0.84 | 19923 | 26048 | 0.76 | 11384 | 23040 | 57.14 | 88.45 | C:97.6%[S:86.1%,D:11.5%],F:0.9%,M:1.5%,n:1614 |
| | TSB (LR/ST2 | 44845 | 34071 | 75.97 | 0.85 | 19241 | 25604 | 0.75 | 11349 | 22722 | 58.98 | 88.74 | C:96.6%[S:83.8%,D:12.8%],F:1.1%,M:2.3%,n:1614 |
| | TSB (SR/LR/ST2) | 48675 | 35380 | 72.69 | 0.83 | 21593 | 27082 | 0.80 | 11833 | 23547 | 54.80 | 86.95 | C:97.9%[S:85.3%,D:12.6%],F:0.5%,M:1.6%,n:1614 |
