## Supplement Table 2 for "Welcome to the big leaves: best practices for improving genome annotation in non-model plant genomes"

*Table S10: Comparison between MAKER and BRAKER*

| | MAKER (RM2+) | | | | | BR (SR) | | | | |
|---|---|---|---|---|---|---|---|---|---|---|
| Species | BUSCO | gFACS | | canonical splice (%) | Gene lengths | BUSCO | gFACS | | canonical splice (%) | Gene lengths |
| | | monoexonic | multiexonic | | | | monoexonic | multiexonic | | |
| ***Arabidopsis thaliana*** | C:90.4%[S:89.0%,D:1.4%],F:3.2%,M:6.4%,n:1614 | 4152 | 18341 | 99.82 | 2342.81 | C:96.3%[S:90.2%,D:6.1%],F:1.0%,M:2.7%,n:1614 | 5066 | 22300 | 99.06 | 2936.07 |
| ***Populus trichocarpa*** | C:19.6%[S:17.9%,D:1.7%],F:1.5%,M:78.9%,n:1614 | 330 | 7273 | 98.39 | 6135.47 | C:97.9%[S:75.0%,D:22.9%],F:1.0%,M:1.1%,n:1614 | 9434 | 38991 | 98.75 | 3142.98 |
| ***Funaria hygrometrica*** | C:79.6%[S:66.7%,D:12.9%],F:4.9%,M:15.5%,n:1614 | 18930 | 25468 | 99.86 | 1868.69 | C:85.8%[S:63.8%,D:22.0%],F:2.9%,M:11.3%,n:1614 | 15084 | 36909 | 99.04 | 2112.89 |

*Table S11: Comparison between Stringtie2 and BRAKER runs*

| Stringtie | | | | | | | | BRAKER | | | | | |
|---|---|---|---|---|---|---|---|---|---|---|---|---|---|
| **Species** | **Run** | **mono** | **multi** | **total** | **ratio** | **BUSCO** | **EnTAP annoation** | **mono** | **multi** | **total** | **ratio** | **BUSCO** | **EnTAP annoation** |
| ***Arabidopsis*** | SR | 6236 | 29287 | 35523 | 0.21 | C:95.5%[S:70.7%,D:24.8%],F:1.1%,M:3.4%,n:1614 | 89.13 | 7515 | 19663 | 27178 | 0.38 | C:95.9%[S:88.2%,D:7.7%],F:0.2%,M:3.9%,n:1614 | 89.87 |
| | LR | 6062 | 30117 | 36179 | 0.20 | C:84.7%[S:61.2%,D:23.5%],F:4.9%,M:10.4%,n:1614 | 83.77 | 7245 | 19128 | 26373 | 0.3787641154 | C:93.9%[S:87.5%,D:6.4%],F:1.1%,M:5.0%,n:1614 | 89.61 |
| | SR+LR | 7266 | 34885 | 42151 | 0.21 | C:93.6%[S:65.1%,D:28.5%],F:0.9%,M:5.5%,n:1614 | 84.52 | 7313 | 19232 | 26545 | 0.3802516639 | C:95.6%[S:88.5%,D:7.1%],F:0.4%,M:4.0%,n:1614 | 89.70 |
| | SR (RM2+) | 6193 | 29287 | 35480 | 0.21 | C:95.5%[S:71.0%,D:24.5%],F:1.1%,M:3.4%,n:1614 | 89.14 | 7656 | 19529 | 27185 | 0.3920323621 | C:95.7%[S:88.8%,D:6.9%],F:0.3%,M:4.0%,n:1614 | 87.94 |
| ***Funaria*** | SR | 13166 | 45543 | 58709 | 0.29 | C:84.5%[S:41.2%,D:43.3%],F:3.0%,M:12.5%,n:1614 | 70.00 | 25184 | 20700 | 45884 | 1.22 | C:84.9%[S:65.1%,D:19.8%],F:2.2%,M:12.9%,n:1614 | 43.27 |
| | SR (RM2+) | 13166 | 45543 | 58709 | 0.29 | C:84.5%[S:41.2%, | 70.00 | 26358 | 20688 | 47046 | 1.274071926 | C:84.7%[S:65.1 | 43.93 |

| **Stringtie** | | | | | | | | **BRAKER** | | | | | |
|---|---|---|---|---|---|---|---|---|---|---|---|---|---|
| | | | | | | D:43.3%],F:3.0%,M:12.5%,n:1614 | | | | | | %,D:19.6%],F:2.2%,M:13.1%,n:1614 | |
| ***Populus*** | SR | 10799 | 38162 | 48961 | 0.28 | C:73.3%[S:45.0%,D:28.3%],F:6.9%,M:19.8%,n:1614 | 82.70 | 13068 | 27773 | 40841 | 0.47 | C:95.6%[S:70.9%,D:24.7%],F:0.8%,M:3.6%,n:1614 | 75.52 |
| | LR | 2744 | 17873 | 20617 | 0.15 | C:51.6%[S:42.1%,D:9.5%],F:12.7%,M:35.7%,n:1614 | 91.17 | 12928 | 25985 | 38913 | 0.50 | C:93.3%[S:72.2%,D:21.1%],F:1.9%,M:4.8%,n:1614 | 76.11 |
| | SR +LR | 3322 | 11067 | 14389 | 0.30 | C:40.3%[S:33.5%,D:6.8%],F:5.3%,M:54.4%,n:1614 | 87.46 | 13014 | 27744 | 40758 | 0.47 | C:94.5%[S:71.0%,D:23.5%],F:1.6%,M:3.9%,n:1614 | 74.97 |
| | SR (RM2+) | 10799 | 38166 | 48965 | 0.28 | C:73.3%[S:45.0%,D:28.3%],F:6.9%,M:19.8%,n:1614 | 82.74 | 14161 | 27932 | 42093 | 0.5069812402 | C:94.6%[S:72.0%,D:22.6%],F:1.7%,M:3.7%,n:1614 | 76.17 |
| ***Liriodendron*** | SR | 16793 | 44714 | 61507 | 0.37 | C:87.1%[S:52.0%,D:35.1%],F:7.2%,M:5.7%,n:1614 | 70.33 | 24180 | 24970 | 49150 | 0.9683620344 | C:81.1%[S:70.9%,D:10.2%],F:7.9%,M:11.0%,n:1614 | 59.67 |
| | LR | 11755 | 21887 | 33642 | 0.54 | C:65.7%[S:52.4%,D:13.3%] | 71.39 | 25137 | 24603 | 49740 | 1.02170467 | C:79.5%[S:69.8%,D:9.7 | 61.40 |

| Stringtie | | | | | | | | BRAKER | | | | | |
|---|---|---|---|---|---|---|---|---|---|---|---|---|---|
| | | | | | | ,F:13.1%, M:21.2%, n:1614 | | | | | | %],F:8.2 %,M:12. 3%,n:16 14 | |
| | SR +LR | 16792 | 44716 | 61508 | 0.37 | C:77.3%[ S:58.5%, D:18.8%] ,F:13.3%, M:9.4%,n :1614 | 70.240141 | 26198 | 25432 | 51630 | 1.03 | C:80.1% [S:69.8 %,D:10. 3%],F:8. 8%,M:11 .1%,n:16 14 | 60.20 |
| | SR (RM2+) | 17018 | 40257 | 57275 | 0.42 | C:75.2%[ S:45.2%, D:30.0%], F:10.0%, M:14.8%, n:1614 | 67.84 | 25815 | 24851 | 50666 | 1.038791 196 | C:80.8% [S:70.6 %,D:10. 2%],F:8. 4%,M:10 .8%,n:16 14 | 60.40 |
| ***Rosa*** | SR | 13382 | 39857 | 53239 | 0.33 | C:97.0%[ S:68.2%, D:28.8%] ,F:0.7%, M:2.3%,n :1614 | 78.81 | 19147 | 25430 | 44577 | 0.752929 6107 | C:95.8% [S:84.0 %,D:11. 8%],F:0. 4%,M:3. 8%,n:16 14 | 68.15 |
| | LR | 19582 | 54372 | 73954 | 0.36 | C:88.7%[ S:57.0%, D:31.7%] ,F:4.4%, M:6.9%,n :1614 | 71.97 | 19241 | 25604 | 44845 | 0.75 | C:94.4% [S:81.5 %,D:12. 9%],F:1. 2%,M:4. 4%,n:16 14 | 55.45 |
| | SR +LR | 28178 | 76552 | 104730 | 0.37 | C:97.2%[ S:49.9%, D:47.3%] ,F:0.8%, M:2.0%,n :1614 | 71.97 | 21593 | 27082 | 48675 | 0.797319 2526 | C:95.8% [S:83.3 %,D:12. 5%],F:0. 7%,M:3. 5%,n:16 14 | 67.17 |

*Table S12: Stringtie proteins- before and after frame-selection*

| | | BUSCO on transdecoder protein | BUSCO on gFACs protein |
|---|---|---|---|
| | SR | C:93.9%[S:67.4%,D:26.5%],F:1.4%,M:4.7%,n:1614 | C:39.8%[S:29.7%,D:10.1%],F:7.5%,M:52.7%,n:1614 |
| | LR | C:85.5%[S:61.3%,D:24.2%],F:5.0%,M:9.5%,n:1614 | C:34.9%[S:27.1%,D:7.8%],F:4.8%,M:60.3%,n:1614 |
| | SR+LR | C:94.5%[S:65.1%,D:29.4%],F:0.4%,M:5.1%,n:1614 | C:40.9%[S:31.2%,D:9.7%],F:4.8%,M:54.3%,n:1614 |
| ***Arabidopsis*** | SR (RM2+) | C:96.4%[S:70.9%,D:25.5%],F:0.8%,M:2.8%,n:1614 | C:41.5%[S:31.9%,D:9.6%],F:5.5%,M:53.0%,n:1614 |
| | SR | C:86.7%[S:41.6%,D:45.1%],F:2.4%,M:10.9%,n:1614 | C:47.1%[S:30.4%,D:16.7%],F:5.9%,M:47.0%,n:1614 |
| ***Funaria*** | SR (RM2+) | C:86.7%[S:41.6%,D:45.1%],F:2.4%,M:10.9%,n:1614 | C:47.1%[S:30.4%,D:16.7%],F:5.9%,M:47.0%,n:1614 |
| | SR | C:95.8%[S:57.2%,D:38.6%],F:1.1%,M:3.1%,n:1614 | C:42.8%[S:30.2%,D:12.6%],F:9.8%,M:47.4%,n:1614 |
| | LR | C:52.3%[S:42.4%,D:9.9%],F:13.0%,M:34.7%,n:1614 | C:22.4%[S:19.6%,D:2.8%],F:7.0%,M:70.6%,n:1614 |
| | SR +LR | C:86.1%[S:65.3%,D:20.8%],F:7.8%,M:6.1%,n:1614 | C:35.0%[S:30.2%,D:4.8%],F:6.7%,M:58.3%,n:1614 |
| ***Populus*** | SR (RM2+) | C:95.8%[S:57.2%,D:38.6%],F:1.1%,M:3.1%,n:1614 | C:42.8%[S:30.2%,D:12.6%],F:9.8%,M:47.4%,n:1614 |
| | SR | C:89.2%[S:52.7%,D:36.5%],F:6.4%,M:4.4%,n:1614 | |
| | LR | C:66.8%[S:53.0%,D:13.8%],F:13.2%,M:20.0%,n:1614 | C:26.4%[S:21.7%,D:4.7%],F:6.6%,M:67.0%,n:1614 |
| | SR +LR | C:79.1%[S:59.5%,D:19.6%],F:12.9%,M:8.0%,n:1614 | C:35.3%[S:29.9%,D:5.4%],F:9.0%,M:55.7%,n:1614 |
| ***Liriodendron*** | SR (RM2+) | C:89.2%[S:52.7%,D:36.5%],F:6.4%,M:4.4%,n:1614 | C:43.6%[S:29.7%,D:13.9%],F:11.7%,M:44.7%,n:1614 |
| | SR | C:97.7%[S:68.0%,D:29.7%],F:0.6%,M:1.7%,n:1614 | C:43.7%[S:33.0%,D:10.7%],F:7.6%,M:48.7%,n:1614 |
| | LR | C:89.7%[S:57.2%,D:32.5%],F:4.3%,M:6.0%,n:1614 | C:36.6%[S:27.4%,D:9.2%],F:7.3%,M:56.1%,n:1614 |
| ***Rosa*** | SR +LR | C:98.0%[S:49.6%,D:48.4%],F:0.4%,M:1.6%,n:1614 | C:47.2%[S:31.8%,D:15.4%],F:9.5%,M:43.3%,n:1614 |

*Table S13: Lirodendron masked and unmasked*

| Masking | Mono | Multi | Input format |
|---|---|---|---|
| Unmasked | 59021 | 148473 | BR (SR) |
| | 45299 | 133575 | BR (SR/LR) |
| | 67597 | 127324 | BR (LR) |
| Repeat masked | 13403 | 38752 | BR (SR) |
| | 12748 | 38259 | BR (SR/LR) |
| | 15568 | 34774 | BR (LR) |
| Additional masking for LTRs | 13566 | 38222 | BR (SR/RM2+) |

*Table S15: Overlaps between BR (SR) and BR (SR/ST2) in Liriodendron*

| Run | Mono | Multi | Overlap with BR (SR) |
|---|---|---|---|
| **BR (SR)** | 13404 | 38755 | |
| **TSB (SR/ST2)** | 24180 | 24970 | 15234 |
| **TSB (SR/OrthoDB)** | 23190 | 23518 | 11111 |
