## Supplement Figures 1 for "Welcome to the big leaves: best practices for improving genome annotation in non-model plant genomes"

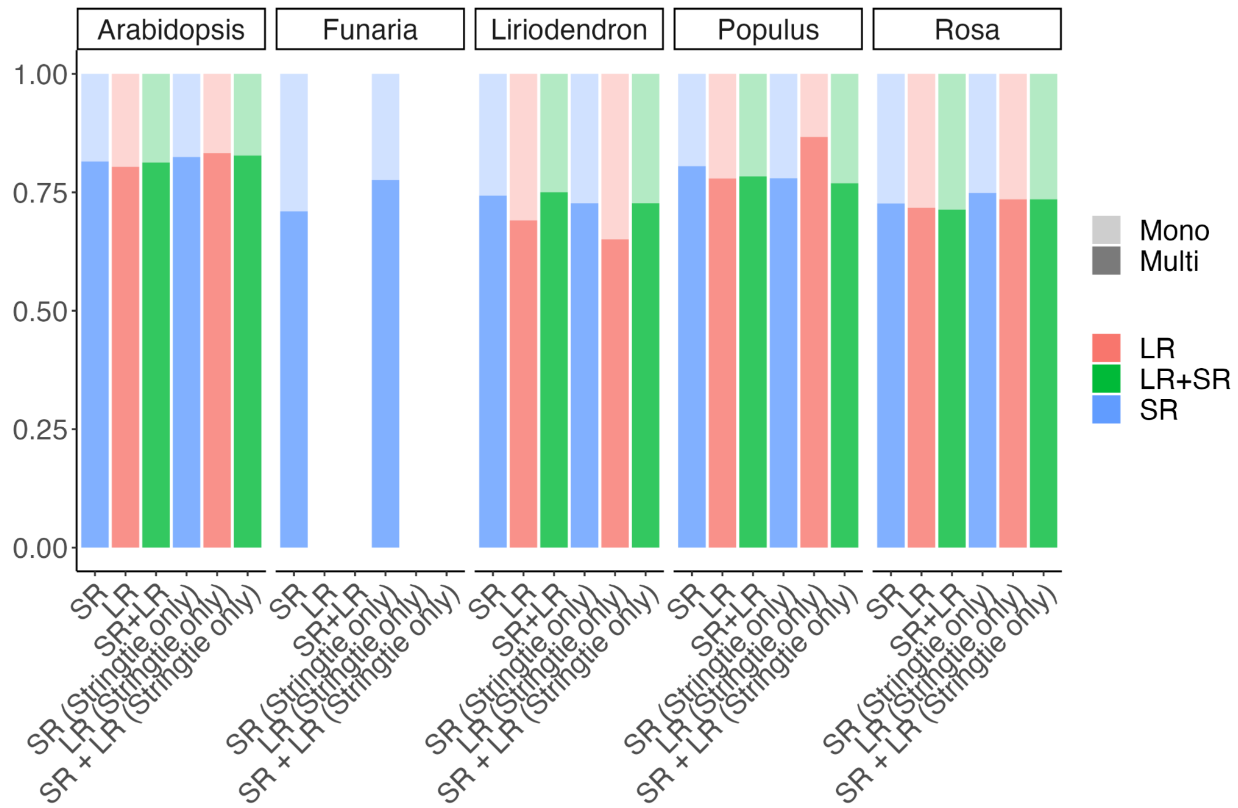


*Figure S1: The mono: multi ratios remain the same regardless of read input type. The lighter shaded area represents the percentage of mono-exonics, and the darker shaded area represents the percentage of multi-exonics. As seen across species, the ratios of mono: multiexonic genes remain similar across species. Comparisons between read-input types may be affected by coverage depth, and type of long read data (PacBio/Nanopore), and that may have affected the ratio.*

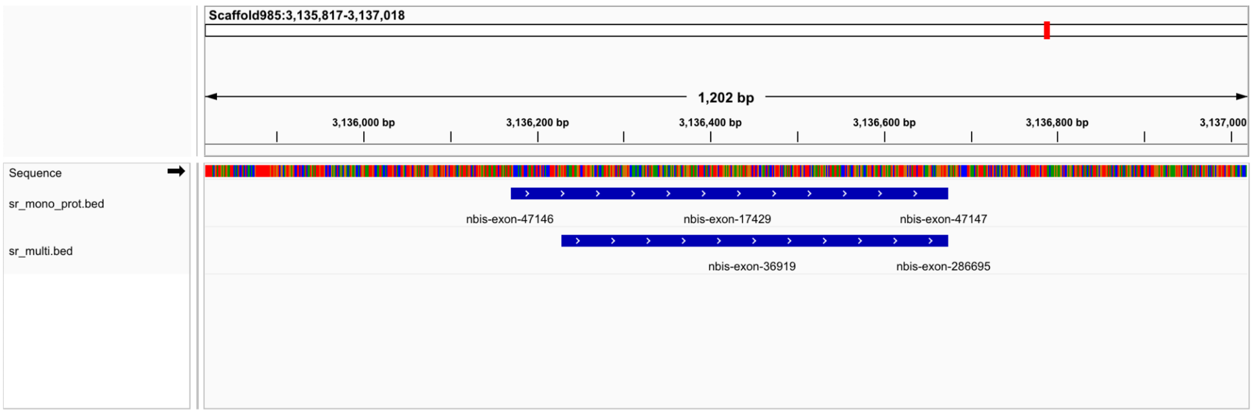


*Figure S2: Genes predicted as mono-exonic from the SR + PROT(ST) run overlapping with multis from the SR run in Liriodendron*
